# Electrical Stimulus Artifact Cancellation and Neural Spike Detection on Large Multi-Electrode Arrays

**DOI:** 10.1101/089912

**Authors:** Gonzalo E. Mena, Lauren E. Grosberg, Sasidhar Madugula, Paweł Hottowy, Alan Litke, John Cunningham, E.J. Chichilnisky, Liam Paninski

## Abstract

Simultaneous electrical stimulation and recording using multi-electrode arrays can provide a valuable technique for studying circuit connectivity and engineering neural interfaces. However, interpreting these measurements is challenging because the spike sorting process (identifying and segregating action potentials arising from different neurons) is greatly complicated by electrical stimulation artifacts across the array, which can exhibit complex and nonlinear waveforms, and overlap temporarily with evoked spikes. Here we develop a scalable algorithm based on a structured Gaussian Process model to estimate the artifact and identify evoked spikes. The effectiveness of our methods is demonstrated in both real and simulated 512-electrode recordings in the peripheral primate retina with single-electrode and several types of multi-electrode stimulation. We establish small error rates in the identification of evoked spikes, with a computational complexity that is compatible with real-time data analysis. This technology may be helpful in the design of future high-resolution sensory prostheses based on tailored stimulation (e.g., retinal prostheses), and for closed-loop neural stimulation at a much larger scale than currently possible.

**Author Summary:** Simultaneous electrical stimulation and recording using multi-electrode arrays can provide a valuable technique for studying circuit connectivity and engineering neural interfaces. However, interpreting these recordings is challenging because the spike sorting process (identifying and segregating action potentials arising from different neurons) is largely stymied by electrical stimulation artifacts across the array, which are typically larger than the signals of interest. We develop a novel computational framework to estimate and subtract away this contaminating artifact, enabling the large-scale analysis of responses of possibly hundreds of cells to tailored stimulation. Importantly, we suggest that this technology may also be helpful for the development of future high-resolution neural prosthetic devices (e.g., retinal prostheses).

## 1 Introduction

Simultaneous electrical stimulation and recording with multi-electrode arrays (MEAs) serves at least two important purposes for investigating neural circuits and for neural engineering. First, it enables the probing of neural circuits, leading to improved understanding of circuit anatomy and function [1–6]. Second, it can be used to assess and optimize the performance of brain-machine interfaces, such as retinal prostheses [7,8], by exploring the patterns of stimulation required to achieve particular patterns of neural activity. However, identifying neural activity in the presence of artifacts introduced by electrical stimulation is a major challenge, and automation is required to efficiently analyze recordings from large-scale MEAs. Furthermore, closed-loop experiments require the ability to assess neural responses to stimulation in real time to actively update the stimulus and probe the circuit, so the automated approach for identifying neural activity must be fast [9,10].

Spike sorting methods [11–13] allow identification of neurons from their spatio-temporal electrical footprints recorded on the MEA. However, these methods fail when used on data corrupted by stimulation artifacts. Although technological advances in stimulation circuitry have enabled recording with significantly reduced artifacts [14–18], identification of neural responses from artifact-corrupted recordings still presents a challenging task — even for human experts — since these artifacts can be much larger than spikes [19], overlap temporally with spikes, and occupy a similar temporal frequency band as spikes.

Although a number of approaches have been previously proposed to tackle this problem [20–23], there are two shortcomings we address here. First, previous approaches are based on restrictive assumptions on the frequency of spikes and their latency distribution (e.g, stimulation-elicited spikes have to occur at least 2ms following stimulus onset). Consequently, it becomes necessary to discard non-negligible portions of the recordings [19,24], leading to biased results that may miss the low-latency regimes where the most interesting neuronal dynamics occur [25,26]. Second, all of these methods have a local nature, i.e., they are based on electrode-wise estimates of the artifact that don’t exploit the shared spatio-temporal information present in MEAs. In general this leads to suboptimal performance. Therefore, a scalable computational infrastructure for spike sorting with stimulation artifacts in large-scale setups is necessary.

This paper presents a method to identify single-unit spike events in electrical stimulation and recording experiments using large-scale MEAs. We develop a modern, large-scale, principled framework for the analysis of neural voltage recordings that have been corrupted by stimulation artifacts. First, we model this highly structured artifact using a structured Gaussian Process (GP) to represent the observed variability across stimulation amplitudes and in the spatial and temporal dimensions measured on the MEA. Next, we introduce a spike detection algorithm that leverages the structure imposed in the GP to achieve a fast and scalable implementation. Importantly, our algorithm exploits many characteristics that make this problem tractable, allowing it to separate the contributions of artifact and neural activity to the observed data. For example, the artifact is smooth in certain dimensions, with spatial footprints that are different than those of spikes. Also, artifact variability is different than that of spikes: while the artifact does not substantially change if the same stimulus is repeated, responses of neurons in many stimulation regimes are stochastic, enhancing identifiability.

The effectiveness of our method is demonstrated by comparison on simulated data and against human-curated inferred spikes extracted from real data recorded in primate retina. Although some features of our method are context-dependent, we discuss extensions to other scenarios, stressing the generality of our approach.

**Fig 1.**
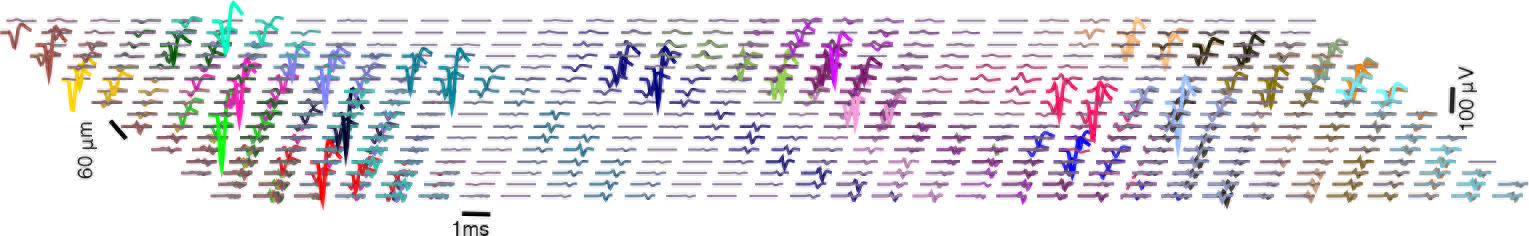
Overlapping electrical images of 24 neurons (different colors) over the MEA, aligned to onset of spiking at *t* = 0.5*ms*. Each trace represents the time course of voltage at a certain electrode. For each neuron, traces are only shown in the electrodes with a strong enough signal. Only a subset of neurons visible on the MEA are shown, for better visibility.

## 2 Materials and Methods

In this section we develop a method for identifying neural activity in response to electrical stimulation. We assume access to voltage recordings *Y*(*e*, *t*, *j*, *i*) in a MEA with *e* = 1,…, *E* electrodes (here, *E* = 512), during *t* = 1,… *T* timepoints (e.g., *T* = 40, corresponding to 2 milliseconds for a 20Khz sampling rate) after the presentation of *j* = 1,…, *J* different stimuli, each of them being a current pulse of increasing amplitudes *a*_*j*_ (in other words, the *a*_*j*_ are magnification factors applied to an unitary pulse). For each of these stimuli *n*_*j*_ trials or repetitions are available; *i* indexes trials. Each recorded data segment is modeled as a sum of the true signal of interest *s* (neural spiking activity on that electrode), plus two types of noise.

The first noise source, *A*, is the large artifact that results from the electrical stimulation at a given electrode. This artifact has a well defined structure but its exact form in any given stimulus condition is not known *a priori* and must be estimated from the data and separated from occurrences of spikes. Although in typical experimental setups one will be concerned with data coming from many different stimulating electrodes, for clarity we start with the case of just a single stimulating electrode; we will generalize this below.

The second source of noise, *ϵ*, is additive spherical Gaussian observation noise; that is, 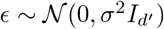, with 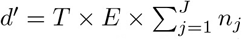. This assumption is rather restrictive and we assume it here for computational ease, but refer the reader to the discussion for a more general formulation that takes into account correlated noise.

Additionally, we assume that *electrical images* (EI) [27,28] — the spatio-temporal collection of action potential shapes on every electrode *e* — are available for all the *N* neurons under study. In detail, each of these EIs are estimates of the voltage deflections produced by a spike over the array in a time window of length *T*′. They are represented as matrices with dimensions *E* × *T*′ and can be obtained in the absence of electrical stimulation, using standard large-scale spike sorting methods (e.g. [12]). Fig 1 shows examples of many EIs, or templates, obtained during a visual stimulation experiment.

Finally, we assume the observed traces are the linear sum of neural activity, artifact, and other noise sources; that is:

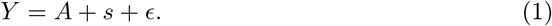

Similar linear decompositions have been recently utilized to tackle related neuroscience problems [12,29].

Figure 2 illustrates the difficulty of this problem: even if 1) for low-amplitude stimuli the artifact may not heavily corrupt the recorded traces and 2) the availability of several trials can enhance identifiability — as traces with spikes and no spikes naturally cluster into different groups — in the general case we will be concerned also with high amplitudes of stimulation. In these regimes, spikes could significantly overlap temporarily with the artifact, and occur with high probability and almost deterministically, i.e., with low latency variability. For example, in the rightmost columns of Figure 2, spike identification is not straightforward since all the traces look alike, and the shape of a typical trace does not necessarily suggest the presence of neural activity. There, inference of neural activity is only possible given a reasonable estimate of the artifact: for instance, under the assumption that the artifact is a smooth function of the stimulus strength, one can make a good initial guess of the artifact by considering the artifact at a lower stimulation amplitude, where spike identification is relatively easier.

**Fig 2.**
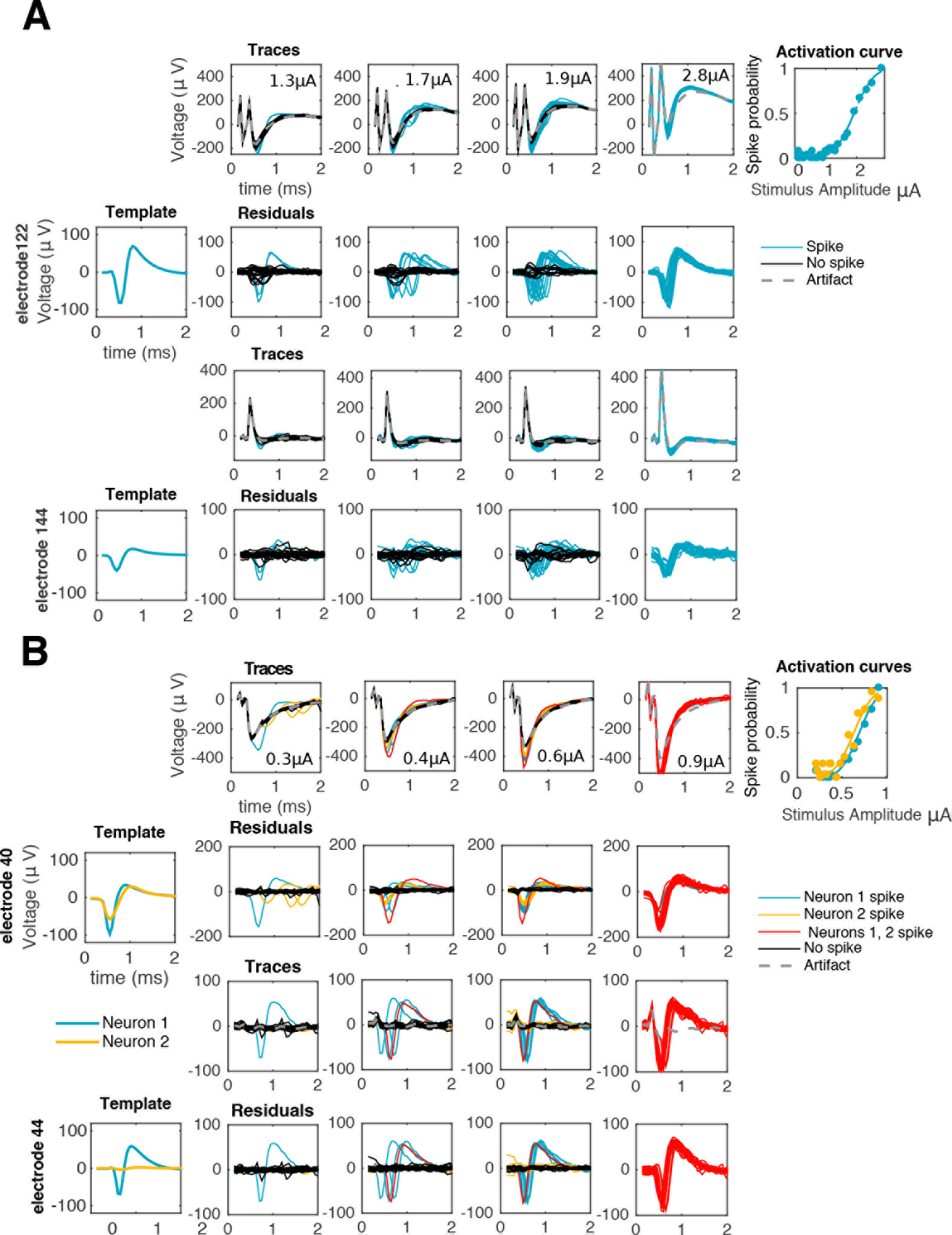
Visual inspection of traces reveals the difficulty of the problem. First column: templates of spiking neurons. Second to fourth columns: responses of one (**A**) or two (**B**) cells to electrical stimulation at increasing stimulation amplitudes as recorded in the stimulating electrode (first rows) or a neighboring, non-stimulating electrode (third rows). If the stimulation artifact is known (gray traces) it can be subtracted from raw traces to produce a baseline (second and fourth rows) amenable for template matching: traces with spike(s) (colored) match, on each electrode, either a translation of a template (**A** and **B**) or the sum of different translations of two or more templates (**B**). As reflected by the activation curves (fifth column) for strong enough stimuli spiking occurs with probability close to one, consistent with the absence of black traces in the rightmost columns.

Therefore, a solution to this problem will rely on a method for an appropriate separation of neural activity and artifact, which in turn requires the use of sensible models that properly capture the structure of the latter; that is, how it varies along the different relevant dimensions. In the following we develop such a method, and divide its exposition in five parts. We start by describing in 2.1 how to model neural activity. Second, in 2.2 we describe the structure of the stimulation artifacts. Third, in 2.3 we propose a GP model to represent this structure. Fourth, in 2.4 we introduce a scalable algorithm that produces an estimate of *A* and *s* given recordings *Y*. Finally, in 2.5 we provide a simplified version of our method and extend it to address multi-electrode stimulation scenarios.

### 2.1 Modeling neural activity

We assume that *s* is the linear superposition of the activities *s*^*n*^ of the *N* neurons involved, i.e. 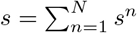. Furthermore, each of these activities is expressed in terms of the binary vectors *b*^*n*^ that indicate spike occurrence and timing: specifically, if 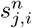 is the neural activity of neuron *n* at trial *i* of the *j*-th stimulation amplitude, we write 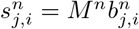, where *M*^*n*^ is a matrix that contains on each row a copy of the EI of neuron *n* (vectorizing over different electrodes) aligned to spiking occurring at different times. Notice that this binary representation immediately entails that: 1) on each trial each neuron fires at most once (this will be the case if we choose analysis time windows that are shorter than the refractory period) and 2) that spikes can only occur over a discrete set of times (a strict subset of the entire recording window), which here corresponds to all the time samples between 0.25 ms and 1.5 ms. We refer the reader to [30] for details on how to relax this simplifying assumption.

### 2.2 Stimulation Artifacts

Electrical stimulation experiments where neural responses are inhibited (e.g., using the neurotoxin TTX) provide qualitative insights about the structure of the stimulation artifact *A*(*e*, *t*, *j*, *i*) (Fig 3); that is, how it varies as a function of all the relevant covariates: space (represented by electrode, *e*), time *t*, amplitude of stimulus *a*_*j*_, and stimulus repetition *i*. Repeating the same stimulation leads to the same artifact, up to small random fluctuations, and so by averaging several trials these fluctuations can be reduced, and we can conceive the artifact as a stack of movies *A*(*e*, *t*, *j*), one for each amplitude of stimulation *a*_*j*_.

We treat the stimulating and non-stimulating electrodes separately because of their observed different qualitative properties.

**Fig. 3.**
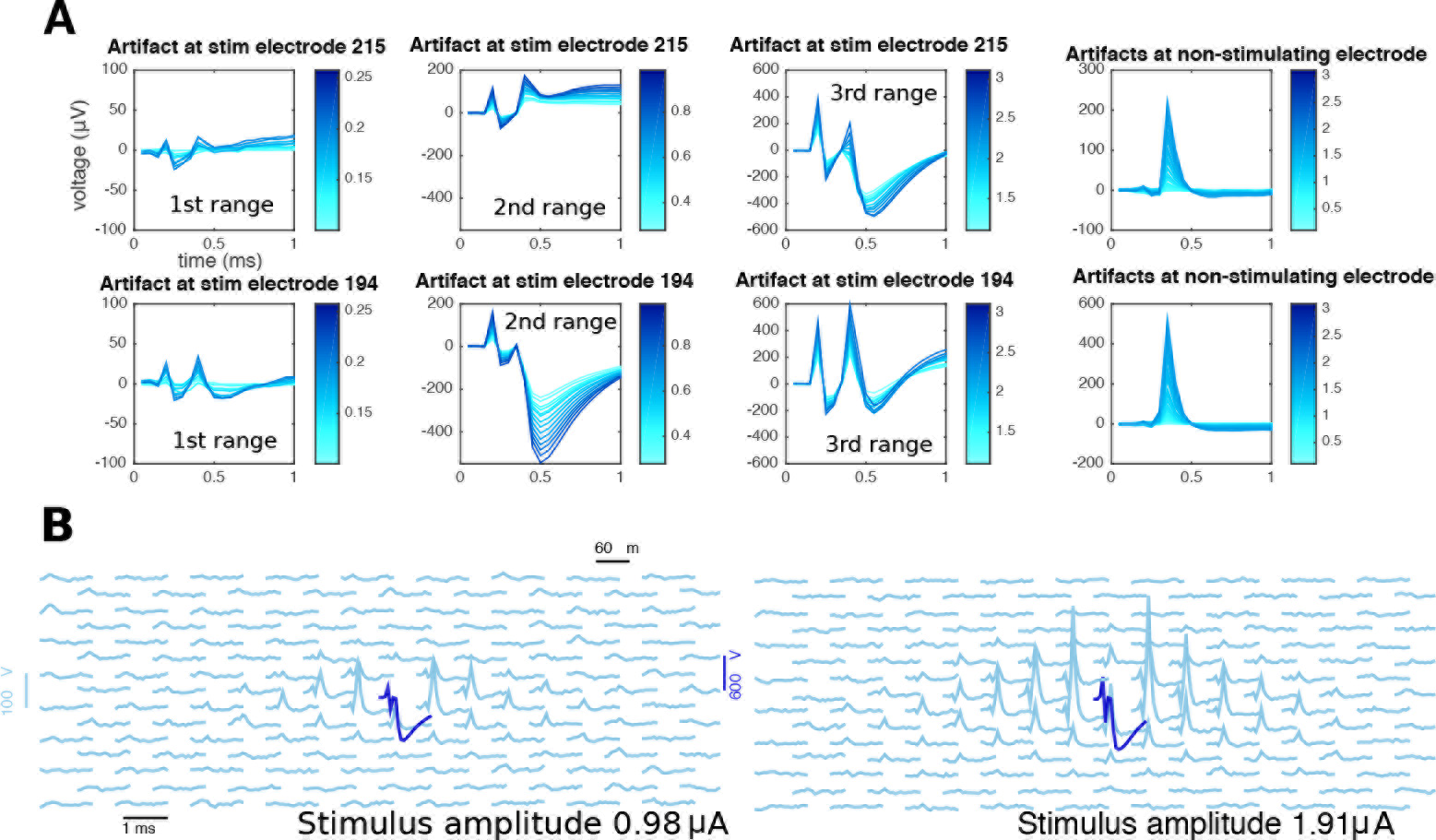
Properties of the electrical stimulation artifact revealed by TTX experiments. (**A**) local, electrode-wise properties of the stimulation artifacts. Overall, magnitude of the artifact increases with stimulation strength (different shades of blue). However, unlike non-stimulating electrodes, where artifacts have a typical shape of a bump around 0.5 ms (fourth column), the case of the stimulating electrode is more complex: besides the apparent increase in artifact strength, the shape itself is not a simple function of stimulating electrode (first and second rows). Also, for a given stimulating electrode the shape of the artifact is a complex function of the stimulation strength, changing smoothly only within certain stimulation ranges: here, responses to the entire stimulation range are divided into three ranges (first, second, and third column) and although traces within each range look alike, traces from different ranges cannot be guessed from other ranges. (**B**) stimulation artifacts in a neighborhood of the stimulating electrode, at two different stimulus strengths (left and right). Each trace represents the time course of voltage at a certain electrode. Notice that stimulating electrode (blue) and non-stimulating electrodes (light blue) are plotted in different scales.

#### 2.2.1 Stimulating electrode

Modeling the artifact in the stimulating electrode requires special care because it is this electrode that typically will capture the strongest neural signal in attempts to directly activate a soma (e.g. Fig 3). The artifact is more complex in the stimulating electrode [16] and has the following properties here: 1) its magnitude is much greater than that of the non-stimulating electrodes; 2) its effect persists at least 2 ms after the onset of the stimulus; and 3) it is a piece-wise smooth, continuous function of the stimulus strength (Fig 3A). Discontinuities occur at a pre-defined set of stimulus amplitudes, the “breakpoints" (known beforehand), resulting from gain settings in the stimulation hardware that must change in order to apply stimuli of different magnitude ranges [16]. Notice that these discontinuities are a rather technical and context-dependent feature that may not necessarily apply to all stimulation systems, unlike the rest of the properties described here.

#### 2.2.2 Non-stimulating electrodes

The artifact here is much more regular and of lower magnitude, and has the following properties (see Fig 3): 1) its magnitude peaks around .4*ms* following the stimulus onset, and then rapidly stabilizes; 2) the artifact magnitude typically decays with distance from the stimulating electrode; 3) the magnitude of the artifact increases with increasing stimulus strength.

Based on these observations, we develop a general framework for artifact modeling based on Gaussian processes.

### 2.3 A structured Gaussian process model for stimulation artifacts

From the above discussion we conclude that the artifact is highly non-linear (on each coordinate), non-stationary (i.e., the variability depends on the value of each coordinate), but structured. The Gaussian process (GP) framework [31] provides powerful and computationally scalable methods for modeling non-linear functions given noisy measurements, and leads to a straightforward implementation of all the usual operations that are relevant for our purposes (e.g. extrapolation and filtering) in terms of some tractable conditional Gaussian distributions.

To better understand the rationale guiding the choice of GPs, consider first a simple Bayesian regression model for the artifact as a noisy linear combination of *B* basis functions Φ_*b*_(*e*, *t*, *j*) (e.g. polynomials); that is, 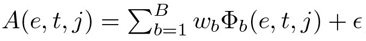, with a regularizing prior *p*(*w*) on the weights. If *p*(*w*) and *ϵ* are modeled as Gaussian, and if we consider the collection of *A*(*e*, *t*, *j*) values (over all electrodes *e*, timesteps *t*, and stimulus amplitude indices *j*) as one large vector *A*, then this translates into an assumption that the vector *A* is drawn from a high-dimensional Gaussian distribution. The prior mean *μ* and covariance *K* of *A* can easily be computed in terms of Φ and *p*(*w*). Importantly, this simple model provides us with tools to estimate the posterior distribution of *A* given partial noisy observations (for example, we could estimate the posterior of *A* at a certain electrode if we are given its values on the rest of the array). Since *A* in this model is a stochastic process (indexed by *e*, *t*, and *j*) with a Gaussian distribution, we say that *A* is modeled as a Gaussian process, and write 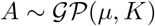.

The main problem with the approach sketched above is that one has to solve some challenging model selection problems: what basis functions Φ_*i*_ should we choose, how large should *M* be, what parameters should we use for the prior *p*(*w*), and so on. We can avoid these issues by instead directly specifying the covariance *K* and mean *μ* (instead of specifying *K* and *μ* indirectly, through *p*(*w*), Φ, etc.).

The parameter *μ* informs us about the mean behavior of the samples from the GP (here, the average values of the artifact). Briefly, we estimate 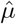 by taking the mean of the recordings at the lowest stimulation amplitude and then subtract off that value from all the traces, so that *μ* can be assumed to be zero in the following. We refer the reader to S1 Text and S1 Fig for details, and stress that all the figures shown in the main text are made after applying this mean-subtraction pre-processing operation.

Next we need to specify *K*. This “kernel" can be thought of as a square matrix of size dim(*A*) × dim(*A*), where dim(*A*) is as large as *T* × *E* × *J* ~ 10^6^ in our context. This number is large enough so all elementary operations (e.g. kernel inversion) are prohibitively slow unless further structure is imposed on *K* — indeed, we need to avoid even storing *K* in memory, and estimating such a high-dimensional object is impossible without some kind of strong regularization. Thus, instead of specifying every single entry of *K* we need to exploit a simpler, lower-dimensional model that is flexible enough to enforce the qualitative structure on *A* that we described in the preceding section.

Specifically, we impose a separable Kronecker product structure on *K*, leading to tractable and scalable inferences [32,33]. This Kronecker product is defined for any two matrices as 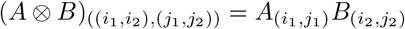. The key point is that this Kronecker structure allows us to break the huge matrix *K* into smaller, more tractable pieces whose properties can be easily specified and matched to the observed data. The result is a much lower-dimensional representation of *K* that serves to strongly regularize our estimate of this very high-dimensional object. In S2 Text we review the main operations from [34] that enable computational speed-ups due to this Kronecker product representation

We state separate Kronecker decompositions for the non-stimulating and stimulating electrodes. For the non-stimulating electrode we assume the following decomposition:

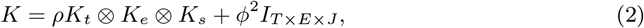

where *K*_*t*_, *K*_*e*_, and *K*_*s*_ are the kernels that account for variations in the time, space, and stimulus magnitude dimensions of the data, respectively. One way to think about the Kronecker product *K*_*t*_ ⊗ *K*_*e*_ ⊗ *K*_*s*_ is as follows: to draw a sample from a GP with mean zero and covariance *K*_*t*_ ⊗ *K*_*e*_ ⊗ *K*_*s*_, start with an array *z*(*t*, *e*, *s*) filled with independent standard normal random variables, then apply independent linear filters in each direction *t*, *e*, and *s* so that the marginal covariances in each direction correspond to *K*_*t*_, *K*_*e*_, and *K*_*s*_, respectively. The dimensionless quantity *ρ* is used to control the overall magnitude of variability and the scaled identity matrix *ϕ*^2^*I*_dim(*A*)_ is included to allow for slight unstructured deviations from the Kronecker structure. Notice that we distinguish between this extra prior variance *ϕ*^2^ and the observation noise variance *σ*^2^, associated with the error term *ϵ* of Eq 1.

Likewise, for the stimulating electrode we consider the kernel:

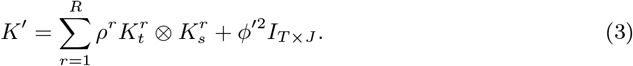

Here, the sum goes over the stimulation ranges defined by consecutive breakpoints; and for each of those ranges, the kernel 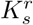 has non-zero off-diagonal entries only for the stimulation values within the *r*-th range between breakpoints. In this way, we ensure artifact information is not shared for stimulus amplitudes across breakpoints. Finally, *ρ*′ and *ϕ*′ play a similar role as in Eq 2.

Now that this structured kernel has been stated it remains to specify parametric families for the elementary kernels *K*_*t*_, *K*_*e*_, *K*_*s*_,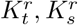. We construct these from the Matérn family, using extra parameters to account for the behaviors described in 2.2.

#### 2.3.1 A non-stationary family of kernels

We consider the Matérn(3/2) kernel, the continuous version of an autoregressive process of order 2. Its (stationary) covariance is given by

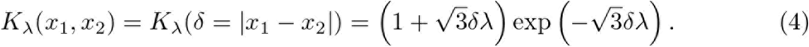

The parameter λ > 0 represents the (inverse) length-scale and determines how fast correlations decay with distance. We use this kernel as a device for representing smoothness; that is, the property that information is shared across a certain dimension (e.g. time). This property is key to induce reasonable extrapolation and filtering estimators, as required by our method (see 2.4). Naturally, given our rationale for choosing this kernel, similar results should be expected if the Matérn(3/2) was replaced by a similar, stationary smoothing kernel.

We induce non-stationarities by considering the family of unnormalized gamma densities *d*_*α*,*β*_(·):

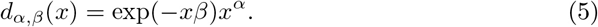

By an appropriate choice of the pair (*α*, *β*) > 0 we aim to expressively represent non-stationary ‘bumps’ in variability. The functions *d*_*α*,*β*_(·) are then used to create a family of non-stationary kernels through the process *Z*_*α*,*β*_ ≡ *Z*_*α*,*β*_ (*x*) = *d*_*α*,*β*_(*x*)*Y*(*x*) where *Y* ~ *GP*(0, *K*_λ_). Thus *Y* here is a smooth stationary process and *d* serves to modulate the amplitude of *Y*. *Z*_*α*,*β*_ is a *bona fide* GP [35] with the following covariance matrix (*D*_*α*,*β*_ is a diagonal matrix with entries *d*_*α*,*β*_(·)):

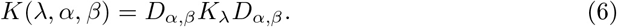

For the non-stimulating electrodes, we choose all three kernels *K*_*t*_, *K*_*e*_, *K*_*s*_ as *K*(λ, *α*, *β*) in Eq 6, with separate parameters λ, *α*, *β* for each. For the time kernels we use time and *t* as the relevant covariate (*δ* in Eq 4 and *x* in Eq 5). The case of the spatial kernel is more involved: although we want to impose spatial smoothness, we also need to express the non-stationarities that depend on the distance between any electrode and the stimulating electrode. We do so by making *δ* represent the distance between recording electrodes, and *x* represent the distance between stimulating and recording electrodes. Finally, for the stimulus kernel we take stimulus strength *a*_*j*_ as the covariate but we only model smoothness through the Matérn kernel and not localization (i.e. *α*, *β* = 0).

Finally, for the stimulating electrode we use the same method for constructing the kernels 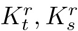 on each range between breakpoints. We provide a notational summary in table 1.

### 2.4 Algorithm

Now we introduce an algorithm for the joint estimation of *A* and *s*, based on the GP model for *A*. Roughly, the algorithm is divided in two stages: first, the hyperparameters that govern the structure of *A* have to be found. This is described in 2.4.1. Second, given the inferred hyperparameters we perform the actual inference of *A*, *s* given these hyperparameters. This is described in 2.4.2 and 2.4.3. We base our approach on posterior inference for *p*(*A*, *s*|*Y*, *θ*, *σ*^2^) ∝ *p*(*Y*|*s*, *A*, *σ*^2^)*p*(*A*|*θ*), where the first factor in the right hand side is the likelihood of the observed data *Y* given *s*, *A*, and the noise variance *σ*^2^, and the second stands for the noise-free artifact prior; *A* ~ *GP*(0, *K*^*θ*^). A summary of all the involved operations is shown in pseudo-code in algorithm 1.

#### 2.4.1 Initialization: hyperparameter estimation

From Eqs (2,3, 4) and 6 the GP model for the artifact is completely specified by the hyperparameters *θ* = (*ρ*, *α*, λ, *β*) and *ϕ*^2^,*ϕ*′^2^. The standard approach for estimating 0 is to optimize the marginal likelihood of the observed data *Y* [31]. However, in this setting computing this marginal likelihood entails summing over all possible spiking patterns s while simultaneously integrating over the high-dimensional vector A; exactly computing this large joint sum and integral is computationally intractable. Instead we introduce a simpler approximation that is computationally relatively cheap and quite effective in practice. We simply optimize the (gaussian) likelihood of *Ã*,

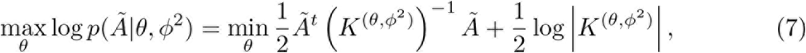

where *Ã* is a computationally cheap proxy for the true *A*. The notation *K*^(*θ*,*ϕ*^2^)^ makes explicit the parametric dependence of the kernels in Eqs 2 and 3, i.e., *K*^(*θ*, *ϕ*^2^)^ = k^*θ*^ + *ϕ*^2^*I*_*T*×*E*×*J*_ with *K*^*θ*^ = *ρK*_*t*_ ⊗ *K*_*e*_ ⊗ *K*_*s*_ for the non-stimulating electrodes (or *K*^(*θ*, *ϕ*′^2^)^ = *K*^*θ*^ + *ϕ*′^2^*I*_*T*×*E*_ and 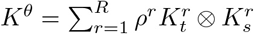 for the stimulating electrode). Due to the Kronecker structure of these matrices, once *Ã* is obtained the terms in Eq. 7 can be computed quite tractably, with computational complexity *O*(*d*^3^), with *d* = max{*E*, *T*, *J*} (max{*T*, *J*} in the stimulating-electrode case), instead of *O*(dim(*A*)^3^), with dim(*A*) = *E* · *T* · *J*, in the case of a general non-structured *K*. Thus the Kronecker assumption here leads to computational efficiency gains of several orders of magnitude. See e.g. [33] for a detailed exposition of efficient algorithmic implementations of all the operations that involve the Kronecker product that we have adopted here; some potential further accelerations are mentioned in the discussion section below.

**Table 1.** Summary of relevant notation.

| Notation | Meaning |
| --- | --- |
| $Y, A, s$ | traces, artifact and neural activity, respectively. |
| $\hat{A}, \hat{s}$ | inferred artifact and neural activity. |
| $t, j, i, e$ | time sample, stimulus index, trial index, electrode index. |
| $T, J, n_j, E$ | number of time samples per recording, number of stimuli, amount of trials per stimulus, number of electrodes in array. |
| $b_{j,i}^n$ | Binary timing vector of spike of neuron $n$, at trial $i$ of $j$-th stimulus. |
| $s_{j,i}^n$ | Action potential (if any) of neuron $n$ at trial $i$ of $j$-th stimulus. |
| $M^n$ | matrix containing action potentials of neuron $n$, aligned to spiking onset at different times as rows. |
| $K_\lambda$ | Matérn(3/2) kernel with inverse length-scale parameter $\lambda$. |
| $K, K'$ | Non-stimulating and stimulating electrodes kernels. |
| $K_t, K_e, K_s$ | time, electrode (space) and stimulus kernels (non-stimulating electrodes). |
| $K_t^r, K_s^r$ | time and stimulus kernels (stimulating electrode) at the $r - th$ range between breakpoints. |
| $R$ | number of intervals between breakpoints. |
| $K_{j,j}$ | sub-matrix of kernel matrix with fixed $j$-th stimulus. |
| $\rho, \rho^r$ | dimensionless factors for stimulating and non-stimulating electrode kernels. |
| $\alpha, \beta$ | parameters of gamma ‘envelope’ $d_{\alpha,\beta}(x) = x^\alpha \exp(-x\beta)$. |
| $\theta$ | vector of kernel hyperparameters: $\theta = (\rho, \alpha, \lambda, \beta)$ and $K^\theta = K_t, K_e, K_s$ (non-stimulating electrodes). |
| $\phi^2$ | noise variance of the artifact. |
| $\sigma^2$ | noise variance of recorded traces. |
| $K^{(\theta, \phi^2)}, K^{(\theta, \phi'^2)}$ | Makes explicit the dependence of $K, K'$ on parameters $K^{(\theta, \phi^2)} = K, K^{(\theta, \phi'^2)} = K'$ |

Now we need to define *Ã*. The stimulating electrode case is a bit more straightforward: we have found that setting *Ã* to the mean or median of *Y* across trials and then solving Eq. 7 leads to reasonable hyperparameter settings. The reason is that can neglect the effect of neural activity on traces, as the artifact *A* is much bigger than the effect of spiking activity s on this electrode, and . We estimate distinct kernels 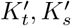 for each stimulating electrode (since from Fig 3A we see that there is a good deal of heterogeneity across electrodes), and each of the ranges between breakpoints. Fig 4B shows an example of some kernels estimated following this approach.

For non-stimulating electrodes, the artifact *A* is more comparable in size to the spiking contributions *s*, and this simple average-over-trials approach was much less successful, explained also by possible corruptions on ‘bad’, broken electrodes which could lead to equally bad hyperparameters estimates. On the other hand, for non-stimulating electrodes the artifact shape is much more reproducible across electrodes, so some averaging over electrodes should be effective. We found that a sensible and more robust estimate can be obtained by assuming that the effect of the artifact is a function of the position relative to the stimulating electrode. Under that assumption we can estimate the artifact by translating, for each of the stimulating electrodes, all the recorded traces as if they had occurred in response to stimulation at the center electrode, and then taking a big average for each electrode. In other words, we estimate

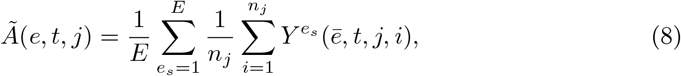

where 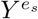 are the traces in response to stimulation on electrode *e*_*s*_ and *ē* is the index of electrode *e* after a translation of electrodes so that *e*_*s*_ is the center electrode. This centered estimate leads to stable values of *θ*, since combining information across many stimulating electrodes serves to average-out stimulating-electrode-specific neural activity and other outliers.

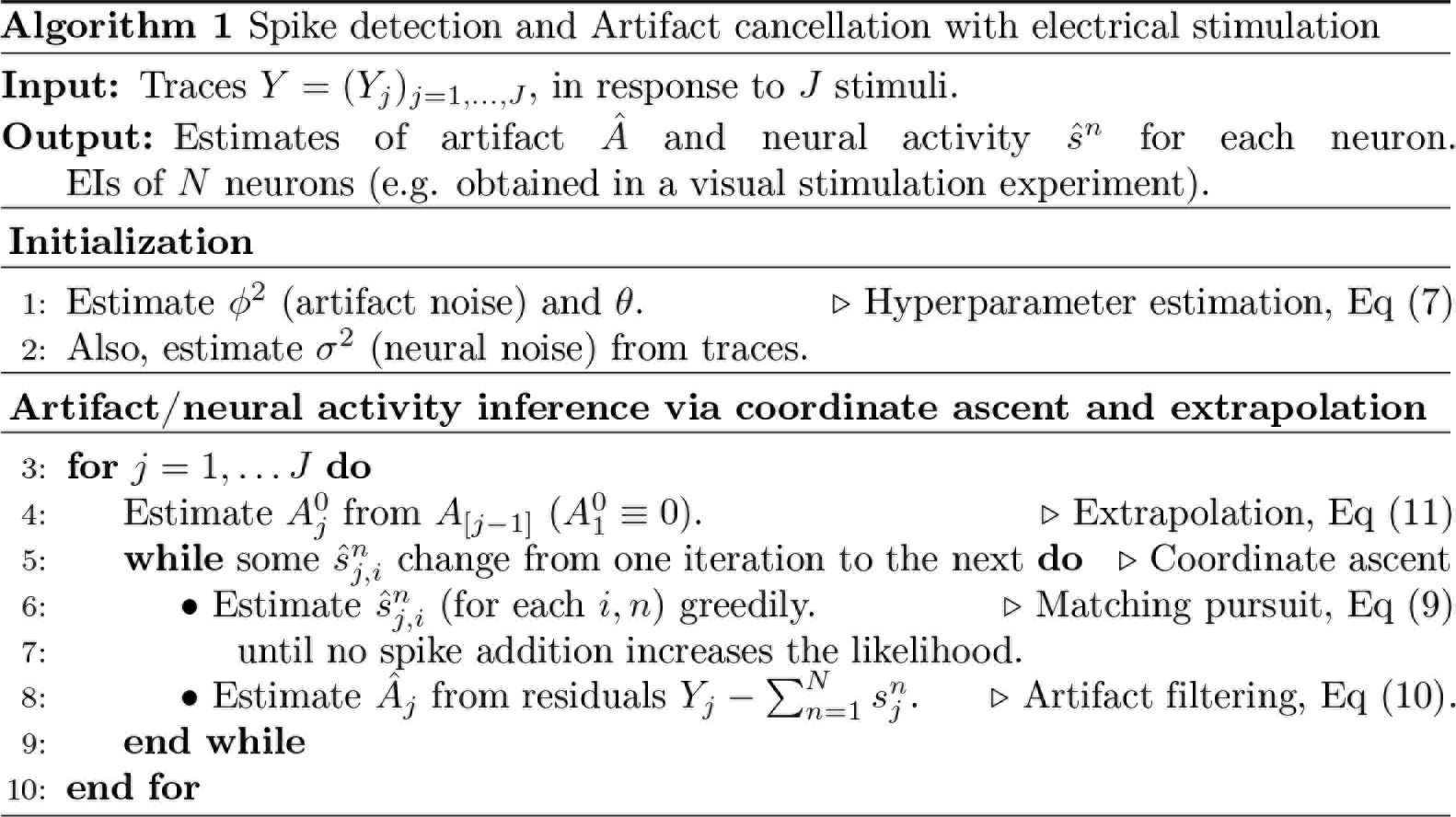

Some implementation details are worth mentioning. First, we do not combine information of all the *E* stimulating electrodes, but rather take a large-enough random sample to ensure the stability of the estimate. We found that using ~ 15 electrodes is sufficient. Second, as the effect of the artifact is very localized in space, we do not utilize all the electrodes, but consider only the ones that are close enough to the center (here, the 25% closest). This leads to computational speed-ups without sacrificing estimate quality; indeed, using the entire array may lead to sub-optimal performance, since distant electrodes essentially contribute noise to this calculation. Third, we do not estimate *ϕ*^2^ by jointly maximizing Eq 7 with respect to (*θ*, *ϕ*). Instead, to avoid numerical instabilities we estimate *ϕ*^2^ directly as the background noise of the fictitious artifact. This can be easily done before solving the optimization problem, by considering the portions of *A* with the lowest artifact magnitude, e.g. the last few time steps at the lowest amplitude of stimulation at electrodes distant from the stimulating electrode. Fig 4A shows an example of kernels *K*_*t*_, *K*_*e*_, and *K*_*s*_ estimated following this approach.

#### 2.4.2 Coordinate Ascent

Once the hyperparameters *θ* are known we focus on the posterior inference for *A*, *s* given *θ* and observed data *Y*. The non-convexity of the set over which the binary vectors *b*^*n*^ are defined makes this problem difficult: many local optima exist in practice and, as a result, for global optimization there may not be a better alternative than to look at a huge number of possible cases. We circumvent this cumbersome global optimization by taking a greedy approach, with two main characteristics: first, joint optimization over *A* and *s* is addressed with alternating ascent (over *A* with *s* held fixed, and then over *s* with *A* held fixed). Alternating ascent is a common approach for related methods in neuroscience (e.g. [12,29]), where the recordings are modeled as an additive sum of spiking, noise, and other terms. Second, data is divided in batches corresponding to the same stimulus amplitude, and the analysis for the (*j* + 1)-th batch starts only after definite estimates *ŝ*_[*j*]_ and *Â*_[*j*]_ have already been produced ([*j*] denotes the set {1,…, *j*}). Moreover, this latter estimate of the artifact is used to initialize the estimate for *A*_*j*+1_ (intuitively, we borrow strength from lower stimulation amplitudes to counteract the more challenging effects of artifacts at higher amplitudes). We address each step of the algorithm in turn below. For simplicity, we describe the details only for the non-stimulating electrodes. Treatment of the stimulating electrode is almost the same but demands a slightly more careful handling that we defer to 2.4.4.

**Fig. 4.**
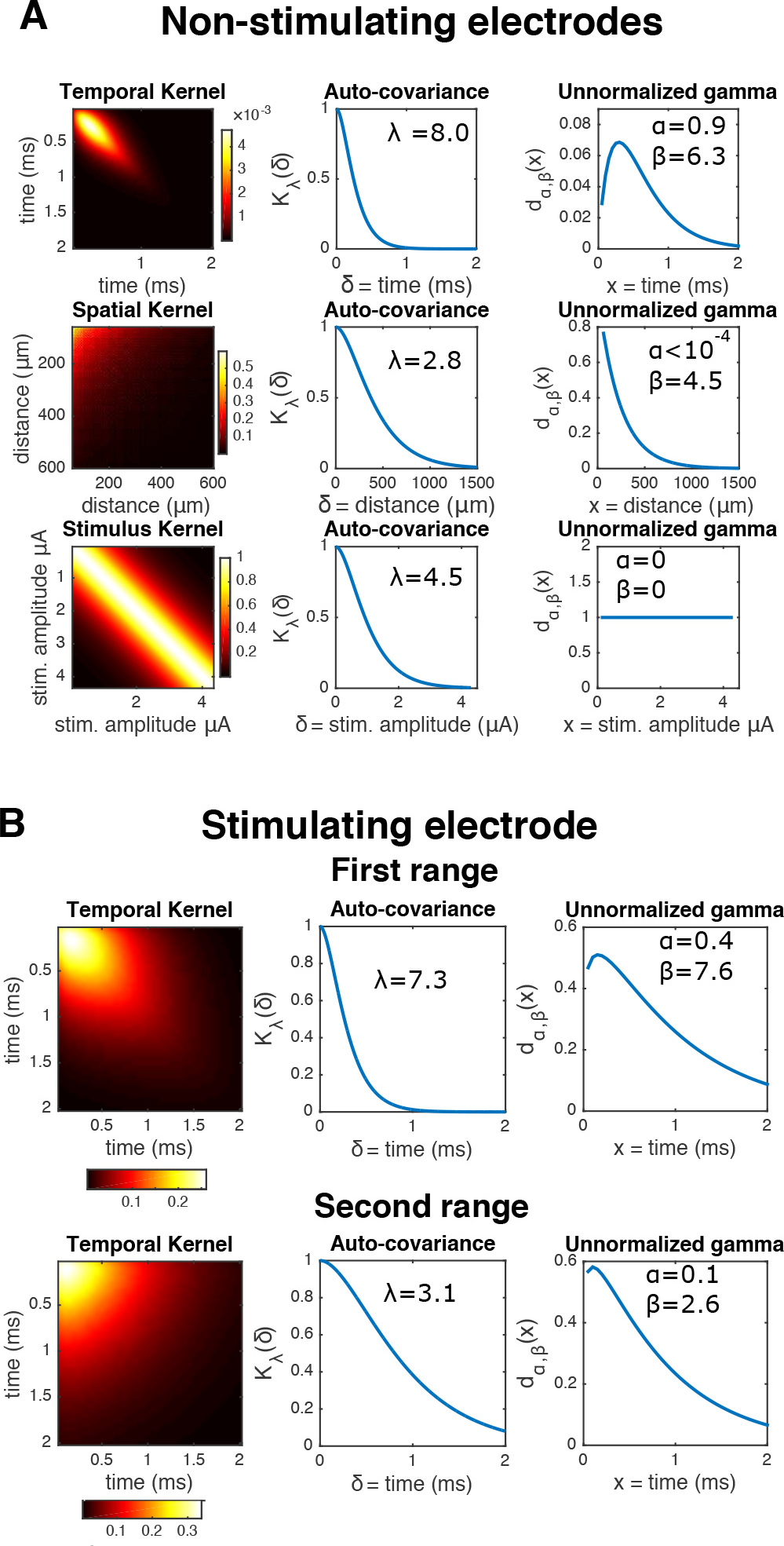
Examples of learned GP kernels. **A** *Left*: inferred kernels *K*_*t*_, *K*_*e*_, *K*_*s*_ in the top, center, and bottom rows, respectively. *Center*: corresponding stationary autocovariances from the Matérn(3/2) kernels (Eq 4). *Right*: corresponding unnormalized ‘gamma-like’ envelopes *d*_*α*,*β*_ (Eq 5). The inferred quantities are in agreement with what is observed in Fig 3B: first, the shape of temporal term *d*_*α*,*β*_ reflects that the artifact starts small, then the variance amplitude peaks at ~ .5 ms, and then decreases rapidly. Likewise, the corresponding spatial *d*_*α*, *β*_ indicates that the artifact variability induced by the stimulation is negligible for electrodes greater than 700 microns away from the stimulating electrode. **B** Same as **A**), but for the stimulating electrode. Only temporal kernels are shown, for two inter-breakpoint ranges (first and second rows, respectively).

Given the batch *Y*_*j*_ and an initial artifact estimate 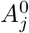 (see 2.4.3) we alternate between neural activity estimation *ŝ*_*j*_ given a current artifact estimate, and artifact estimation *Â*_*j*_ given the current estimate of neural activity. This alternating optimization stops when changes in every 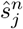 are sufficiently small, or nonexistent.

**Matching pursuit for neural activity inference.** Given the current artifact estimate *Â*_*j*_ we maximize the conditional distribution for neural activity 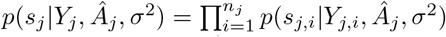, which corresponds to the following sparse regression problem (the set *S* embodies our constraints on spike occurrence and timing):

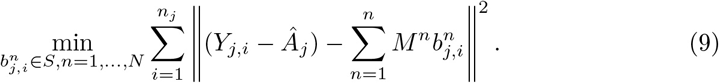

We seek to find the allocation of spikes that will lead the best match with the residuals (*Y*_*j*, *i*_ − *Â*_*j*_). We follow a standard template-matching-pursuit greedy approach (e.g. [12]) to locally optimize Eq 9: specifically, for each trial we iteratively search for the best choice of neuron/time, then subtract the corresponding neural activity until the proposed updates no longer lead to increases in the likelihood.

**Filtering for artifact inference.** Given the current estimate of neural activity *ŝ*_*j*_ we maximize the posterior distribution of the artifact, that is, 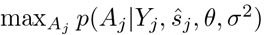, which here leads to the posterior mean estimator (again, the overline indicates mean across the *n*_*j*_ trials):

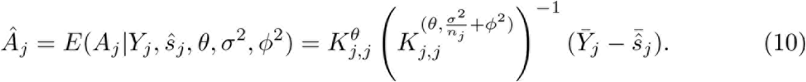

This operation can be understood as the application of a linear filter. Indeed, by appealing to the eigendecomposition of 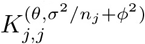 we see this operator shrinks the *m*-th eigencomponent of the artifact by a factor of *κ*_*m*_/(*κ*_*m*_ + *σ*^2^/*n*_*j*_ + *ϕ*^2^) (*κ*_*m*_ is the *m*-th eigenvalue of 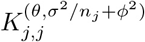), exerting its greatest influence where *κ*_*m*_ is small. Notice that in the extreme case that *σ*^2^/*n*_*j*_ + *ϕ*^2^ is very small compared to the *κ*_*m*_ then 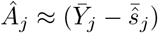, i.e., the filtered artifact converges to the simple mean of spike-subtracted traces.

**Convergence.** Remarkably, in practice often only a few (e.g. 3) iterations of coordinate ascent (neural activity inference and artifact inference) are required to converge to a stable solution 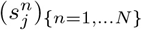. The required number of iterations can vary slightly, depending e.g. on the number of neurons or the signal-to-noise; i.e., EI strength versus noise variance.

#### 2.4.3 Iteration over batches and artifact extrapolation

The procedure described in 2.4.2 is repeated in a loop that iterates through the batches corresponding to different stimulus strengths, from the lowest to the highest. Also, when doing *j* → *j* + 1 an initial estimate for the artifact 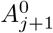 is generated by extrapolating from the current, faithful, estimate of the artifact up to the *j*-th batch. This extrapolation is easily implemented as the mean of the noise-free posterior distribution in this GP setup, that is:

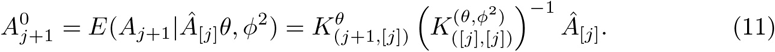

Importantly, in practice this initial estimate ends up being extremely useful, as in the absence of a good initial estimate, coordinate ascent often leads to poor optima. The very accurate initializations from extrapolation estimates help to avoid these poor local optima (see Fig 8).

We note that both for the extrapolation and filtering stages we still profit from the scalability properties that arise from the Kronecker decomposition. Indeed, the two required operations — inversion of the kernel and the product between that inverse and the vectorized artifact — reduce to elementary operations that only involve the kernels *K*_*e*_, *K*_*t*_, *K*_*s*_ [33].

#### 2.4.4 Integrating the stimulating and non-stimulating electrodes

Notice that the same algorithm can be implemented for the stimulating electrode, or for all electrodes simultaneously, by considering equivalent extrapolation, filtering, and matched pursuit operations. The only caveat is that extrapolation across stimulation amplitude breakpoints does not make sense for the stimulating electrode, and therefore, information from the stimulating electrode must not be taken into account at the first amplitude following a breakpoint, at least for the first matching pursuit-artifact filtering iteration.

#### 2.4.5 Further computational remarks

Note the different computational complexities of artifact related operations (filtering, extrapolation) and neural activity inference: while the former depends (cubically) only on *T, E, J*, the latter depends (linearly) on the number of trials *n*_*j*_, the number of neurons, and the number of electrodes on which each neuron’s EI is significantly nonzero. In the data analyzed here, we found that the fixed computational cost of artifact inference is typically bigger than the per-trial cost of neural activity inference. Therefore, if spike sorting is required for big volumes of data (*n*_*j*_ ⪢ 1) it is a sensible choice to avoid unnecessary artifact-related operations: as artifact estimates are stable after a moderate number of trials (e.g. *n*_*j*_ = 50), one could estimate the artifact with that number, subtract that artifact from traces and perform matching pursuit for the remaining trials. That would also be helpful to avoid unnecessary multiple iterations of the artifact inference - spike inference loop.

### 2.5 Simplifications and extensions

#### 2.5.1 A simplified method

We now describe a way to reduce some of the computations associated with algorithm 1. This simplified method is based on two observations: first, as discussed above, if many repetitions are available, the sample mean of spike subtracted traces over trials should already provide an accurate artifact estimator, making filtering (Eq 10) superfluous. (Alternatively, one could also consider the more robust median over trials; in the experiments analyzed here we did not find any substantial improvement with the median estimator.) Second, as artifact changes smoothly across stimulus amplitudes, it is reasonable to use the artifact estimated at condition *j* as an initialization for the artifact estimate at the (*j* + 1)th amplitude. Naturally, if two amplitudes are too far apart this estimator breaks down, but if not, it circumvents the need to appeal to Eq 11.

Thus, we propose a simplified method in which Eq 10 is replaced by the spike-subtracted mean voltage (i.e. skip the filtering step in line 9 of algorithm 1), and Eq 11 is replaced by simple ‘naive’ extrapolation (i.e. avoid kernel-based extrapolation in line 5 of algorithm 1 and just initialize 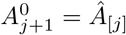). We can derive this simplified estimator as a limiting special case within our GP framework: first, avoiding the filtering operator is achieved by neglecting the noise variances *σ*^2^ and *ϕ*^2^, as this essentially means that our observations are noise-free; hence, there is no need for smoothing. Also, our naive extrapolation proposal can be obtained using an artifact covariance kernel based on Brownian motion in *j* [36].

Finally, notice that the simplified method does not require a costly initialization (i.e, we can skip the maximization of Eq 7 in line 2 of algorithm 1).

#### 2.5.2 Beyond single-electrode stimulation

So far we have focused our attention on single electrode stimulation. A natural question is whether or not our method can be extended to analyze responses to simultaneous stimulation at several electrodes, which is of particular importance for the use of patterned stimulation as a means of achieving selective activation of neurons [28,37]. One simple approach is to simply restrict attention to experimental designs in which the relative amplitudes of the stimuli delivered on each electrode are held fixed, while we vary the overall amplitude. This reduces to a one-dimensional problem (since we are varying just a single overall amplitude scalar). We can apply the approach described above with no modifications to this case, just replacing “stimulus amplitude" in the single-electrode setting with “overall amplitude scale" in the multiple-electrode case.

In this work we consider three types of multiple electrode stimulation: *Bipolar* stimulation, *Local Return* stimulation and *Arbitrary* stimulation patterns. Bipolar stimuli were applied on two neighboring electrodes, and consisted of simultaneous pulses with opposite amplitudes. The purpose was to modulate the direction of the applied electric field [38]. The local return stimulus had the same central electrode current, with simultaneous current waveforms of opposite sign and one sixth amplitude on the six immediately surrounding return electrodes. The purpose of the local return stimulus configuration was to restrict the current spread of the stimulation pulse by using local grounding. More generally, arbitrary stimulation patterns (up to four electrodes) were similarly designed to shape the resulting electric field, and consisted of simultaneous pulses of varied amplitudes.

## 3 Results

We start by showing, in Figure 5, an example of the estimation of the artifact *A* and spiking activity *s* from single observed trials *Y*. Here, looking at individual responses to stimulation provides little information about the presence of spikes, even if the EIs are known. Thus, the estimation process relies heavily on the use of shared information across dimensions: in this example, a good estimate of the artifact was obtained by using information from stimulation at lower amplitudes, and from several trials.

**Fig. 5.**
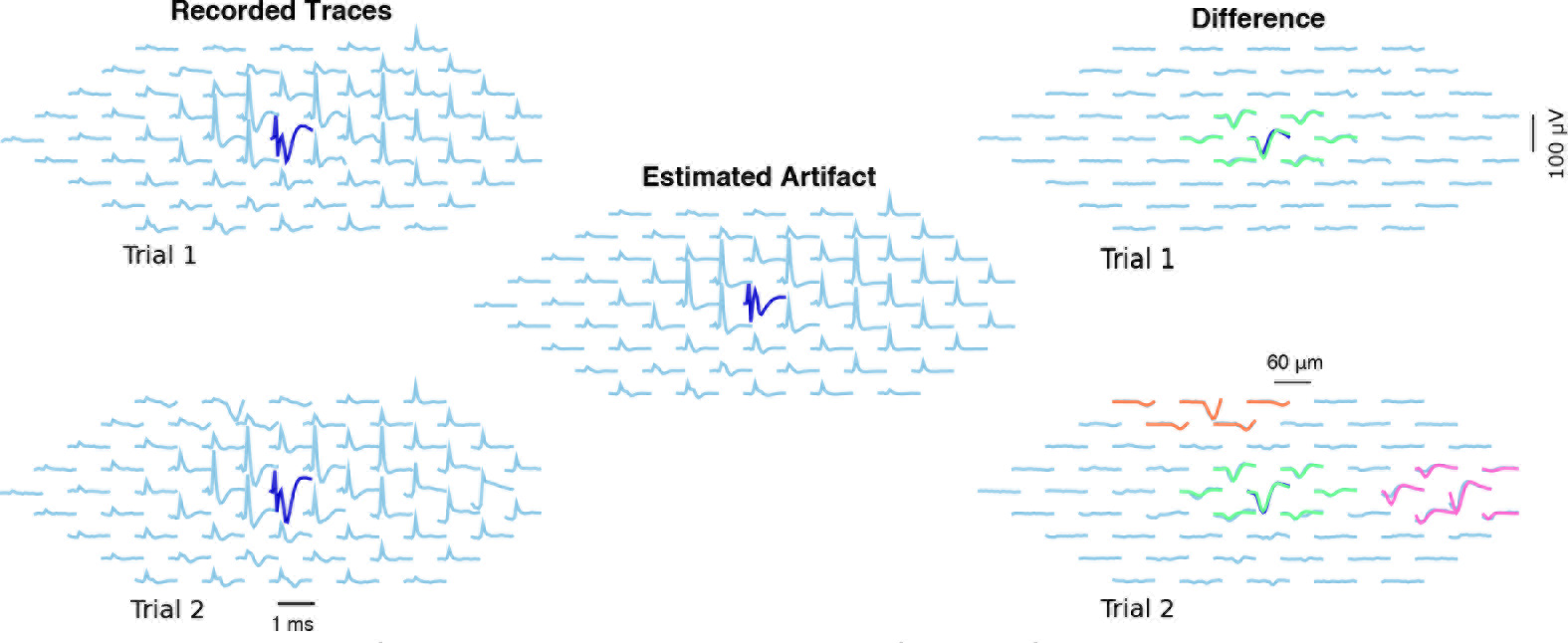
Example of neural activity and artifact inference in a neighborhood of the stimulating electrode. *Left:* Two recordings in response to a 2.01 *μA* stimulus. *Center:* estimated artifact (as the stimulus doesn’t change, it is the same for both trials). *Right:* Difference between raw traces and estimated artifact, with inferred spikes in color. In the first trial (above) one spiking neuron was detected, while in trial 2 (below) three spiking neurons were detected. The algorithm separates the artifact *A* and spiking activity *s* effectively here.

### 3.1 Algorithm validation

We validated the algorithm by measuring its performance both on a large dataset with available human-curated spike sorting and with ground-truth simulated data (we avoid the term ground-truth in the real data to acknowledge the possibility that the human makes mistakes).

#### 3.1.1 Comparison to human annotation

The efficacy of the algorithm was first demonstrated by comparison to human-curated results from the peripheral primate retina. The algorithm was applied to 4,045 sets of traces in response to increasing stimuli. We refer to each of these sets as an *amplitude series*. These amplitude series came from the four stimulation categories described in section 2: single-electrode, bipolar, local return, and arbitrary.

We first assessed the agreement between algorithm and human annotation on a trial-by-trial basis, by comparing the presence or absence of spikes, and their latencies. Results of this trial-by-trial analysis for the kernel-based estimator are shown in Fig 6A. Overall, the results are satisfactory, with an error rate of 0.45%. Errors were the result of either false positives (misidentified spikes over the cases of no spiking) or false negatives (failures in detecting truly existing spikes), whose rates were 0.43% (FPR, false positives over total positives) and 1.08% (FNR, false negatives over total negatives), respectively. For reference, we considered the baseline given by the simple estimator introduced in [20]: there, the artifact is estimated as the simple mean of traces. False negative rates were an order of magnitude larger for the reference estimator, 49% (see S2 Fig for details). In 4.2 we further discuss why this reference method fails in this data.

We observed comparable error rates for the simplified and kernel-based estimator (again, see S2 Fig for details). To further investigate differences in performance, we considered three ‘perturbations’ to real data (restricting our attention to single-electrode stimulation, for simplicity): sub-sampling of trials (by limiting the maximum number of trials per stimulus to 20, 10, 5, and 2), sub-sampling of amplitudes (considering only every other or every other other stimulus amplitude in the sequence), and noise injection, by adding uncorrelated Gaussian noise with standard deviation *σ* = 5,10, or 20*μ* V (this noise adds to the actual noise in recordings that here we estimated below *σ* = 6*μ*V, by using traces in response to low amplitude stimuli far from the stimulation site). Representative results are shown in Fig 6B (but see S3 Fig for full comparisons), and indicate that indeed the kernel-based estimator delivers superior performance in these more challenging scenarios. Thus unless otherwise noted below we focus on results of the full kernel-based estimator, not the simplified estimator; see 3.1.2 and 4.1 for more comparisons between both estimators, and for a broader discussion.

**Fig. 6.**
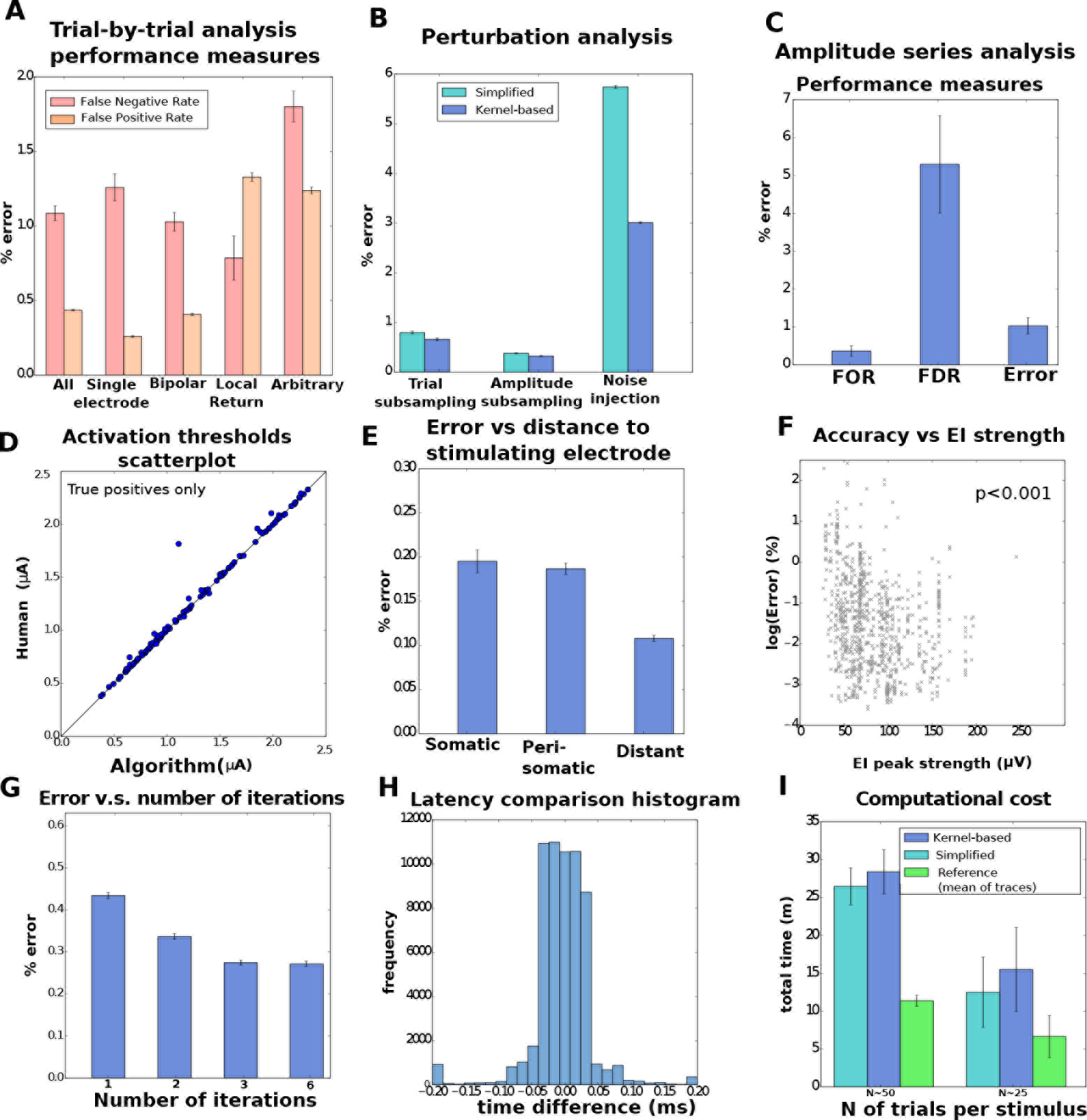
Population results from thirteen retinal preparations reveal the efficacy of the algorithm. **A.** Trial-by-trial wise performance of estimators broken down by the the four types of stimulation considered (total number of trials 1,713,233, see Table 1 S1 text for details). **B.** Trial-by-trial wise performance of estimators to perturbations of real data (only single-electrode): five trials per stimulus for trial subsampling, every other stimulus for amplitude subsampling and *σ* = 20 for noise injection. **C,D.** Amplitude-series wise performance of estimators. C: false omission rate (FOR = FN/(FN+TP)), false discovery rate (FDR = FP/(FP+TP)), and error rate based on the 4,045 available amplitude series (see Table 2 S1 Text for details); **D**: comparison of activation thresholds (human vs. kernel-based algorithm). **E.** Performance measures (trial-by-trial) broken down by distance between neuron and stimulating electrode. **F.** Trial-by-trial error as a function of EI peak strength across all electrodes (only kernel-based). A Spearman correlation test revealed a significant negative correlation. **G.** Error as a function of number of iterations in the algorithm. **H.** For the true positives, histogram of the differences of latencies between human and algorithm. **I.** Computational cost comparison of the three methods for the analysis of single-electrode scans, with 20 to 25 (left) or 50 (right) trials per stimulus.

We also quantified accuracy at the level of the entire amplitude series, instead of individual trials: given an amplitude series we conclude that neural activation is present if the sigmoidal activation function fit (specifically, the CDF of a normal distribution) to the empirical activation curves — the proportion of trials where spikes occurred as a function of stimulation amplitude — exceeds 50% within the ranges of stimulation. In the positive cases, we define the stimulation threshold as the current needed to elicit spiking with 0.5 probability. This number provides an informative univariate summary of the activation curve itself. The obtained results are again satisfactory (Fig 6C). Also, in the case of correctly detected events we compared the activation thresholds (Fig 6D) and found little discrepancy between human and algorithm (with the exception of a single point, which can be better considered as an additional false positive, as the algorithm predicts activation at much smaller amplitude of stimulus; data not shown).

We investigated various covariates that could modulate performance: distance between targeted neuron and stimulating electrode (Fig 6E), strength of the neural signals (Fig 6F) and maximum permitted number of iterations of the coordinate ascent step (Fig 6G). Regarding the first, we divided data by somatic stimulation (stimulating electrode is the closest to the soma), peri-somatic stimulation (stimulating electrode neighbors the closest to the soma) and distant stimulation (neither somatic nor peri-somatic). As expected, accuracies were the lowest when the neural soma was close to the stimulating electrode (somatic stimulation), presumably a consequence of artifacts of larger magnitude in that case. Regarding the second, we found that error significantly decreases with strength of the EI, indicating that our algorithm benefits from strong neural signals. With respect to the third, we observe some benefit from increasing the maximum number of iterations, and that accuracies stabilize after a certain value (e.g. three), indicating that either the algorithm converged or that further coordinate iterations did not lead to improvements.

Finally, we report two other relevant metrics: first, differences between real and inferred latencies (Fig 6H, only for correctly identified spikes) revealed that in the vast majority of cases (>95%) spike times inferred by human vs. algorithm differed by less than 0.1 ms. Second, we assessed computational expenses by measuring the algorithm’s running time for the analysis of a single-electrode scan; i.e, the totality of the 512 amplitude series, one for each stimulating electrode (Fig 6I). The analysis was done in parallel, with twenty threads analyzing single amplitude series (details in S1 Text). We conclude that we can analyze a complete experiment in ten to thirty minutes and that the parallel implementation is compatible with the time scales required by closed-loop pipelines. We further comment on this in 4.3. Comparisons in Fig 6I also illustrate that our methods are 2x-3x slower than the (much less accurate) reference estimator, but that differences between kernel-based and the simplified estimator are rather moderate. This suggests that filtering and extrapolation are inexpensive in comparison to the time spent in the matching pursuit stage of the algorithm, and that the cost of finding the hyper-parameters (only once) is negligible at the scale of the analysis of several hundreds of amplitude series.

We refer the reader to S1 Text for details on population statistics of the analyzed data, exclusion criteria, and computational implementation.

**Fig 7.**
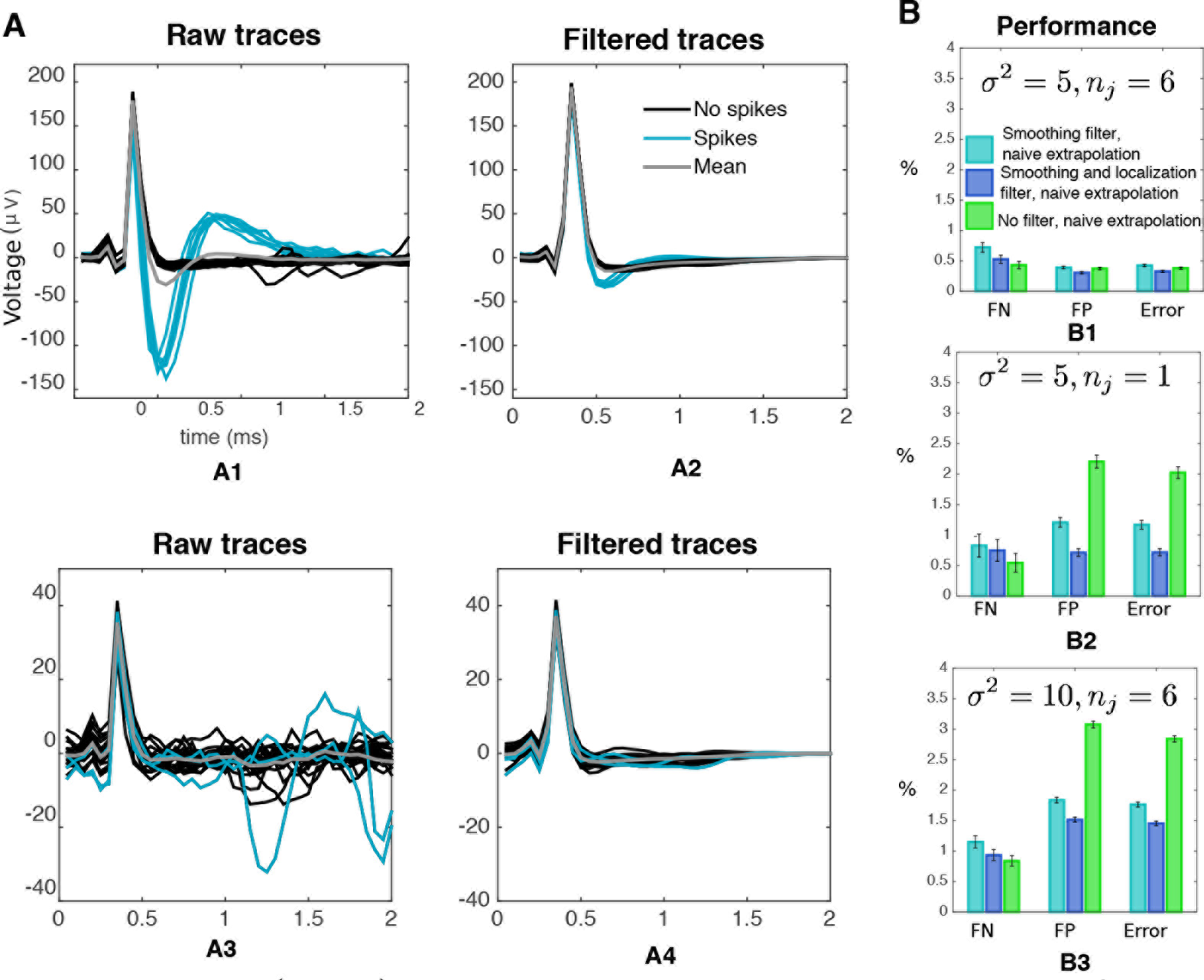
Filtering (Eq 10) leads to a better, less spike-corrupted artifact estimate in our simulations. **A** effect of filtering on traces for two non-stimulating electrodes, at a fixed amplitude of stimulation (2.2*μ*A). *A1,A3* raw traces, *A2,A4* filtered traces. Notice the two main features of the filter: first, it principally affects traces containing spikes, a consequence of the localized nature of the kernel in Eq 2. Second, it helps eliminate high-frequency noise. **B** through simulations, we showed that filtering leads to improved results in challenging situations. Two filters — only smoothing and localization + smoothing — were compared to the omission of filtering. In all cases, to rule out that performance changes were due to the extrapolation estimator, extrapolation was done with the naive estimator. *B1* results in a less challenging situation. *B2* results in the heavily subsampled (*n*_*j*_ = 1) case. *B3* results in the high-noise variance (*σ*^2^ = 10) case.

#### 3.1.2 Simulations

Synthetic datasets were generated by adding artifacts measured in TTX recordings (not contaminated by neural activity *s*), real templates, and white noise, in an attempt to faithfully match basic statistics of neural activity in response to electrical stimuli, i.e., the frequency of spiking and latency distribution as a function of distance between stimulating electrode and neurons (see S5 Fig). These simulations (only on single-electrode stimulation) were aimed to further investigate the differences between the naive and kernel-based estimators, by determining when — and to which extent — filtering (Eq 10) and extrapolation (Eq 11) were beneficial to enhance performance. To address this question, we evaluated separately the effects of the omission and/or simplification of the filtering operation (Eq 10), and of the replacement of the kernel-based extrapolation (Eq 11) by the naive extrapolation estimator that guesses the artifact at the *j*-th amplitude of stimulation simply as the artifact at the *j* − 1 amplitude of stimulation.

**Fig 8.**
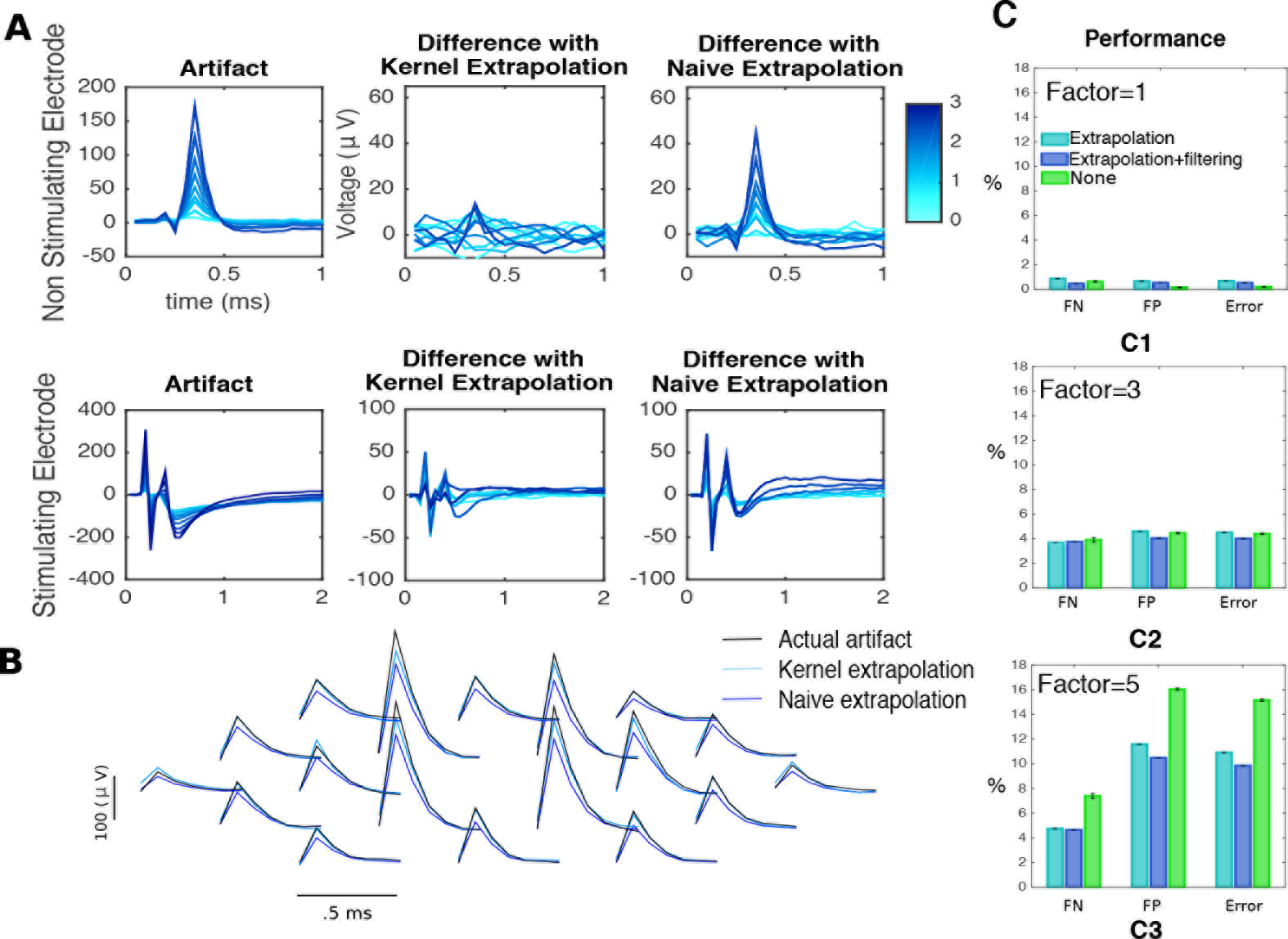
Kernel-based extrapolation (Eq 11) leads to more accurate initial estimates of the artifact. **A** comparison between kernel-based extrapolation and the naive estimator, the artifact at the previous amplitude of stimulation. For a non-stimulating (first row) and the stimulating (second row) electrode, left: artifacts at different stimulus strengths (shades of blue), center: differences with extrapolation estimator (Eq 11), right: differences with the naive estimator. **B** comparison between the true artifact (black), the naive estimator (blue) and the kernel-based estimator (light blue) for a fixed amplitude of stimulus (3.1*μ*A) on a neighborhood of the stimulating electrode (not shown). **C** Through simulations we showed that extrapolation leads to improved results in a challenging situation. Kernel-based extrapolation was compared to naive extrapolation. *C1* results in a less challenging situation. *C2*-*C3* results in the case where the artifact is multiplied by a factor of 3 and 5, respectively.

As the number of trials *n*_*j*_ goes to infinity, or as the noise level *σ* goes to zero, the influence of the likelihood grows compared to the GP prior, and the filtering operator converges to the identity (see Eq 10). However, applied on individual traces, where the influence of this operator is maximal, filtering removes high frequency noise components and variations occurring where the localization kernels do not concentrate their mass (Fig 4A), which usually correspond to spikes. Therefore, in this case filtering should lead to less spike-contaminated artifact estimates. Fig 7B confirms this intuition with results from simulated data: in cases of high *σ*^2^ and small *n*_*j*_ the filtering estimator led to improved results. Moreover, a simplified filter that only consisted of smoothing kernels (i.e. for all the spatial, temporal and amplitude-wise kernels the localization terms *d*_*α*,*β*_ in Eq 5 were set equal to 1, leading to the Matérn kernel in Eq 4) led to more modest improvements, suggesting that the localization terms (Eq 5) — and not only the smoothing kernels — act as sensible and helpful modeling choices.

Likewise, we expect that kernel-based extrapolation leads to improved performance if the artifact magnitude is large compared to the size of the EIs: in this case, differences between the naive estimator and the actual artifact would be large enough that many spikes would be misidentified or missed. However, since kernel-based extrapolation produces better artifact estimates (see Fig 8A-B), the occurrence of those failures should be diminished. Indeed, Fig 8C shows that better results are attained when the size of the artifact is multiplied by a constant factor (or equivalently, neglecting the noise term σ^2^, when the size of the Els is divided by a constant factor). Moreover, the differential results obtained when including the filtering stage suggest that the two effects are non-redundant: filtering and extrapolation both lead to improvements and the improvements due to each operation are not replaced by the other.

**Fig 9.**
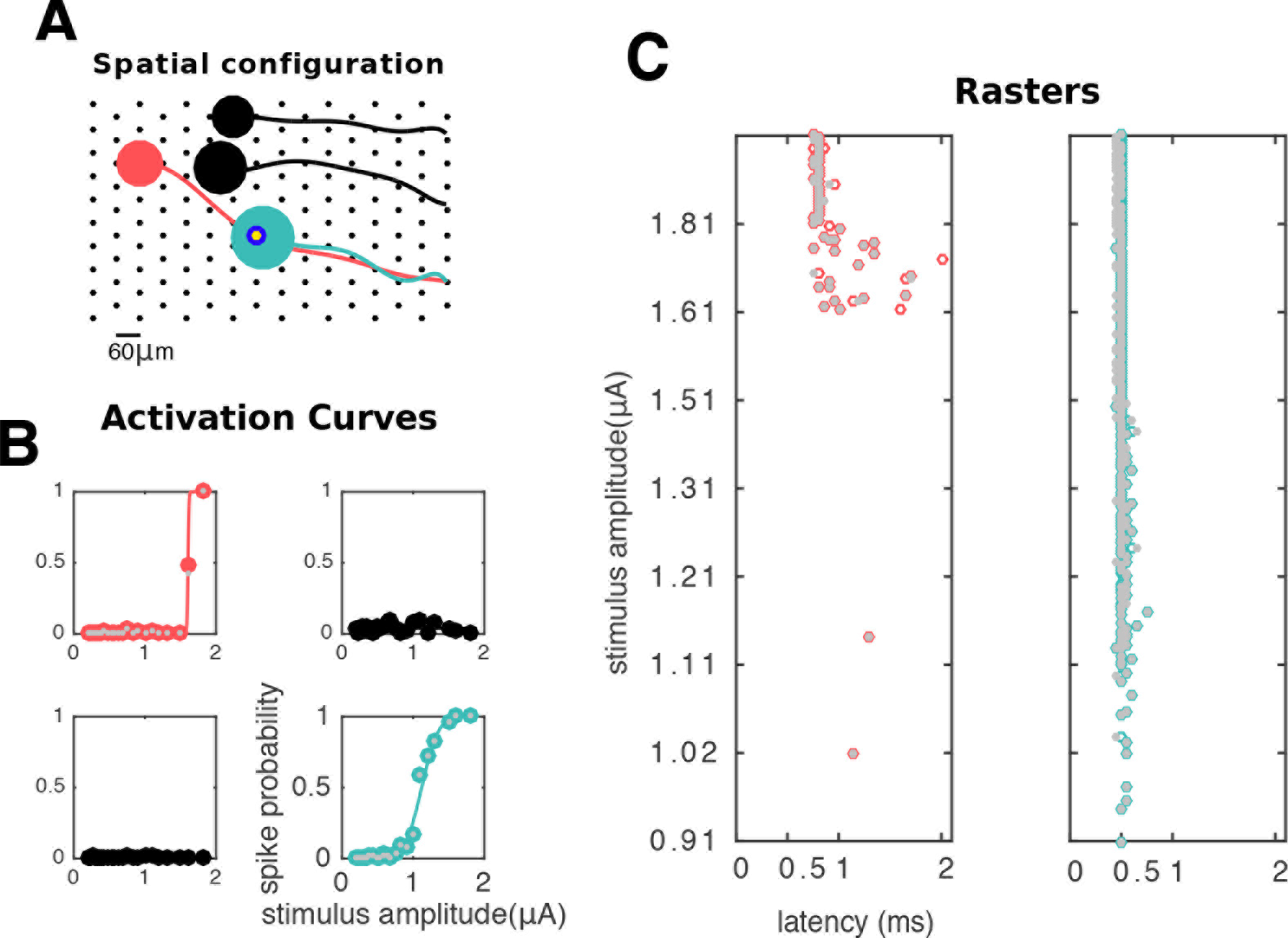
Analysis of responses of neurons in a neighborhood of the stimulating electrode. **A** Spatial configuration: stimulating electrode (blue/yellow annulus) and four neurons on its vicinity. Soma of green neuron and axon of pink neuron overlap with stimulating electrode. **B** Activation curves (solid lines) along with human-curated and algorithm inferred spike probabilities (gray and colored circles, respectively) of all the four cells. Stimulation elicited activation of green and pink neurons; however, the two other neurons remained inactive. **C** Raster plots for the activated cells, with responses sorted by stimulation strength in the y axis. Human and algorithm inferred latencies are in good agreement (gray and colored circles, respectively). Here, direct somatic activation of the green neuron leads to lower-latency and lower-threshold activation than of the pink neuron, which is activated through its axon.

### 3.2 Applications: high resolution neural prosthesis

A prominent application of our method relates to the development of high-resolution neural prostheses (particularly, epi-retinal prosthesis), whose success will rely on the ability to elicit arbitrary patterns of neural activity through the selective activation of individual neurons in real-time [28,39,[40]. For achieving such selective activation in a closed-loop setup, we need to know how different stimulating electrodes activate nearby neurons, information that is easily summarized by the activation curves, with the activation thresholds themselves as proxies. Unfortunately, obtaining this information in real time — as required for prosthetic devices — is currently not feasible since estimation of thresholds requires the analysis of individual responses to stimuli. In 4.3 we discuss in detail how, within our framework, to overcome the stringent time limitations required for such purposes.

**Fig 10.**
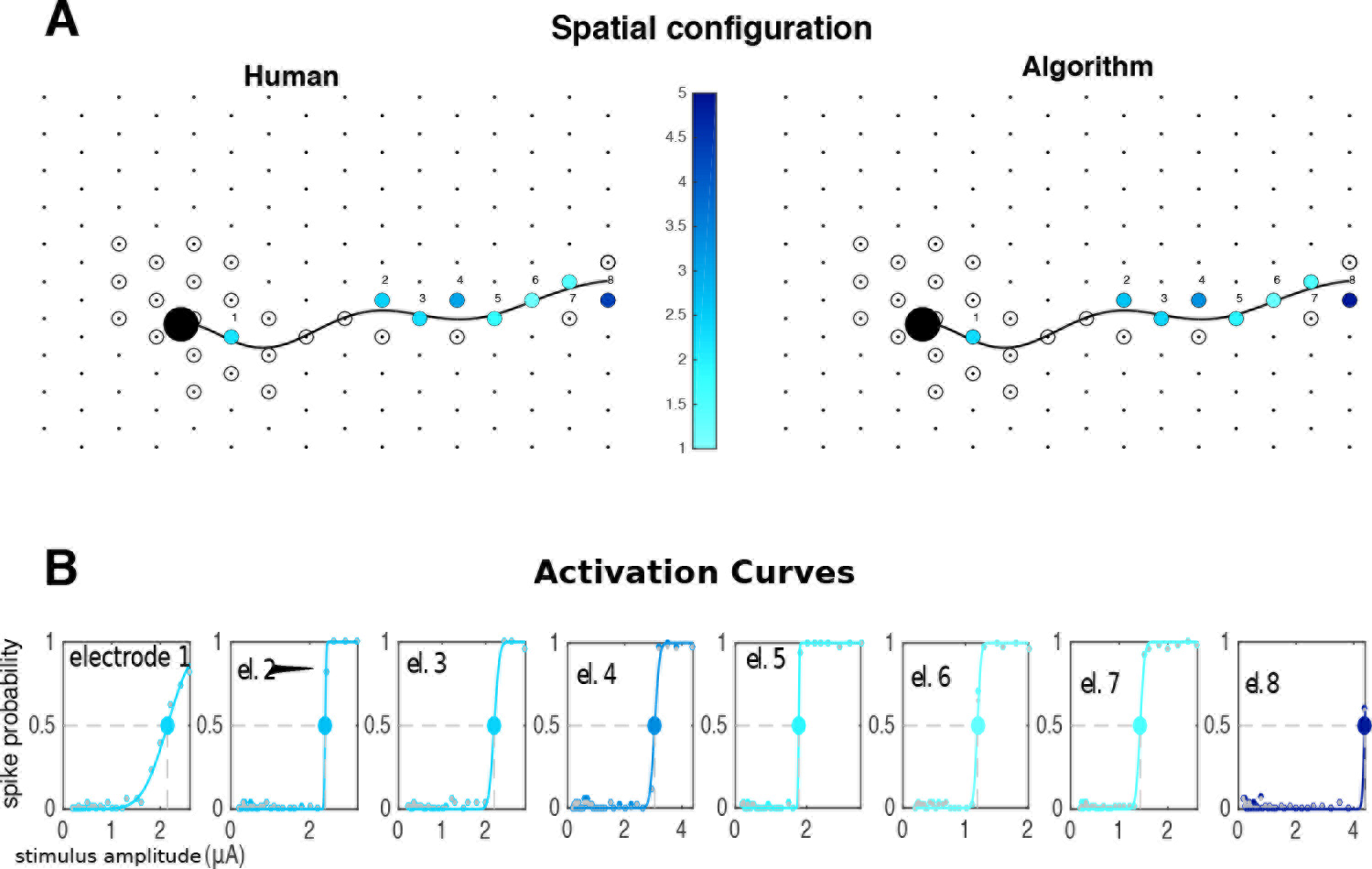
Electrical receptive field of a neuron. **A** spatial representation of the soma (black circle) and axon (black line) over the array. Electrodes where stimulation was attempted are represented by circles, with colors indicating the activation threshold in the case of a successful activation of the neuron within the stimulation range. **B** For those cases, activation curves (solid lines) are shown along with with human and algorithm inferred spike frequencies (gray and colored circles, respectively). Large circles indicate the activation thresholds represented in **A.** In this case, much of the activity is elicited through axonal stimulation, as there is a single electrode close to the soma that can activate the neuron. Human and algorithm are in good agreement.

Figures 9, 10, 11, and 12 show pictorial representations of different features of the results obtained with the algorithm, and their comparison with human annotation. Axonal reconstructions from all of the neurons in the figures were achieved through a polynomial fit to the neuron’s spatial EI, with soma size depending on the EI strength (see [28] for details). Each of these figures provides particular insights to inform and guide the large-scale closed-loop control of the neural population. Importantly, generation of these maps took only minutes on a personal computer, compared to many human hours, indicating feasibility for clinical applications and substantial value for analysis of laboratory experiments [28,40].

Figure 9 focuses on the stimulating electrode’s point of view: given stimulation in one electrode, it is of interest to understand which neurons will get activated within the stimulation range, and how selective that activation can be made. This information is provided by the activation curves, i.e, their steepness and their associated stimulation thresholds. Additionally, latencies can be informative about the spatial arrangement of the system under study, and the mode of neural activation: in this example, one cell is activated through direct stimulation of the soma, and the other, more distant cell is activated through the indirect and antidromic propagation of current through the axon [41]. This is confirmed by the observed latency pattern.

**Fig 11.**
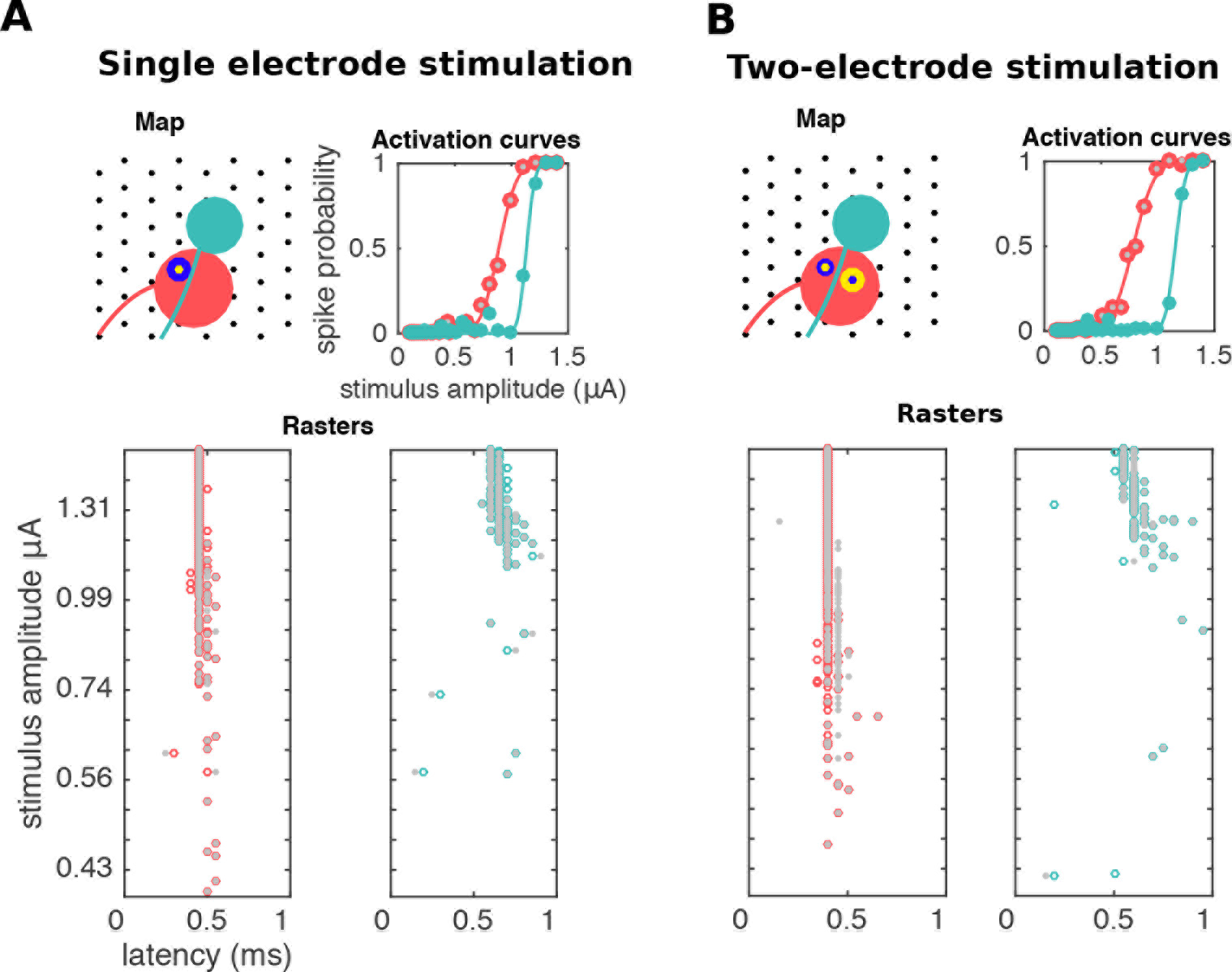
Analysis of differential responses to single (A) and two-electrode (B) stimulation. Gray and colored dots indicate human and algorithm inferences, respectively. In both cases activation of the two neurons is achieved. However, shape of activation curves is modulated by the presence of a current with the same strength and opposite polarity in a neighboring electrode (yellow/blue annulus in **B**): indeed, in this case bipolar stimulation leads to an enhanced ability to activate the pink neuron without activating the green neuron. The algorithm is faithfully able to recover the relevant activation thresholds.

Figure 10 depicts the converse view, focusing on the neuron. Here we aim to determine the cell’s electrical receptive field [37,42] to single-electrode stimulation; that is, the set of electrodes that are able to elicit activation, and in the positive cases, the corresponding stimulation thresholds. These fields are crucial for tailoring stimuli that selectively activate sub-populations of neurons.

Figure 11 shows how the algorithm enables the analysis of responses to bipolar stimulation. This strategy has been suggested to enhance selectivity [43], by differentially shifting the stimulation thresholds of the cells so the range of currents that lead to activation of a single cell is widened. More generally, multi-electrode spatial stimulation patterns have the potential to enhance selectivity by producing an electric field optimized for activating one cell more strongly than others [28], and Fig 11 is a depiction of how our algorithm permits an accurate assessment of this potential enhancement.

Finally, Fig 12 shows a large-scale summary of the responses to single-electrode stimulation. There, a population of ON and OFF parasol cells was stimulated at many different electrodes close to their somas, and each of those cells was then labeled by the lowest achieved activation threshold. These maps provide a proxy of the ability to activate cells with single-electrode stimulation, and of the different degrees of difficulty in achieving activation. Since in many cases only as few as 20% of the neurons can be activated [44], the information about which cells were activated can provide a useful guide for the on-line development of more complex multiple electrode stimulation patterns that activate the remaining cells.

**Fig. 12.**
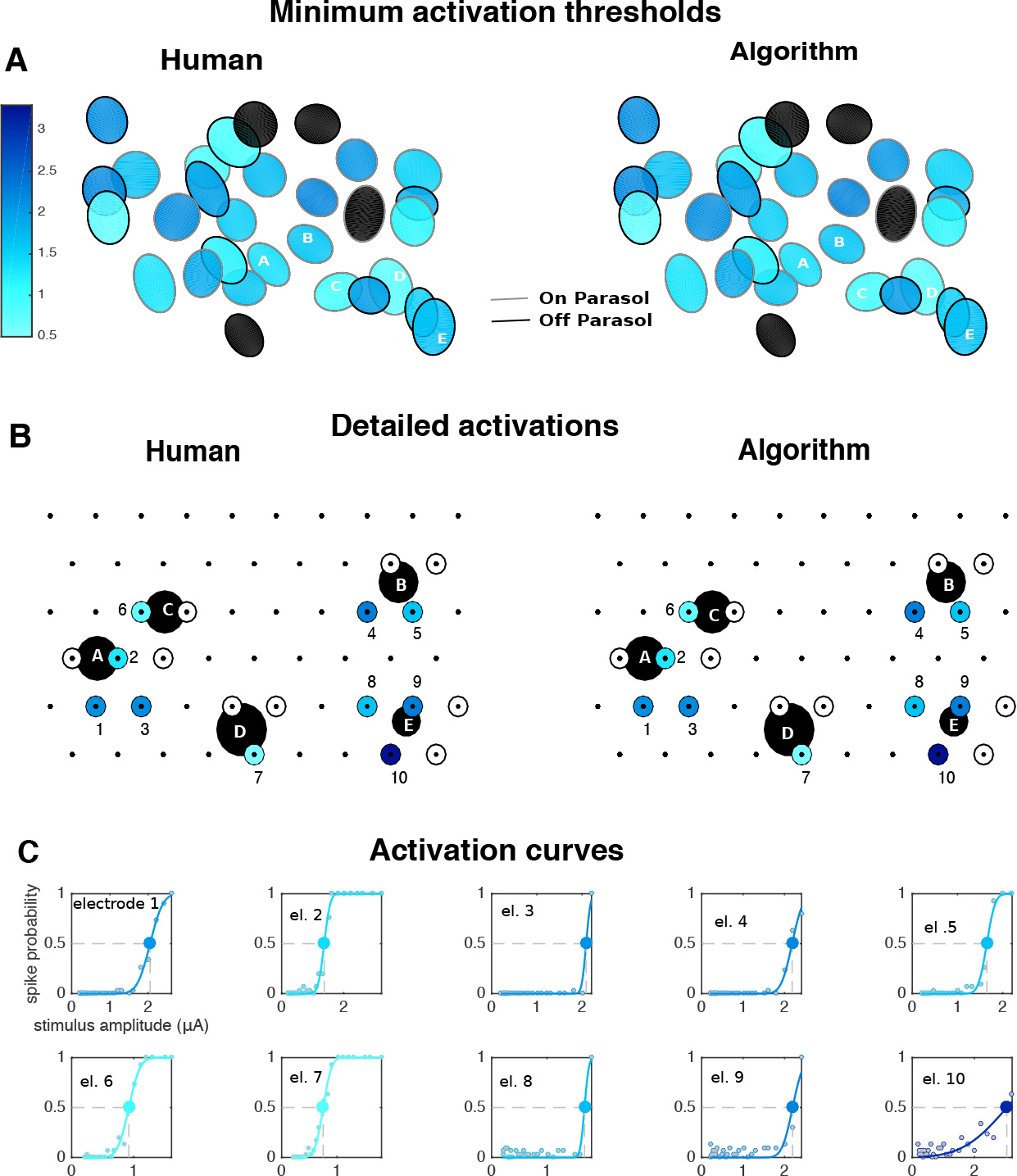
Large-scale analysis of the stimulation of a population of parasol cells. For each neuron, one or more stimulating electrodes in a neighborhood of neural soma were chosen for stimulation. **A** Receptive fields colored by the lowest achieved stimulation threshold (black if activation was not achieved). **B** Inferred somas (big black circles) of the neurons labeled A-E in **A**), showing which electrodes were chosen for stimulation (small circles) and whether activation was achieved (colors). **C** Activation curves (solid lines) of the neurons in **B** for the successful activation cases. Gray and colored dots represent human and algorithm results, respectively, and large circle indicates stimulation thresholds.

## 4 Discussion

Now we discuss the main features of the algorithm in light of the results and sketch some extensions to enable the analysis of data in contexts that go beyond those analyzed here.

### 4.1 Simplified vs. full kernel-based estimators

Figures 6B, 7B, 8C, and S3 Fig illustrate some cases where the full kernel-based estimator outperforms the simplified artifact estimator. These cases correspond to heavy sub-sampling or small signal-to-noise ratios, where the data do not adequately constrain simple estimators of the artifact and the full Bayesian approach can exploit the structure in the problem to obtain significant improvements. In closed-loop experiments (discussed below in 4.3) experimental time is limited, and the ability to analyze fewer trials without loss of accuracy opens up the possibility for new experimental designs that may not have been otherwise feasible. That said, it is useful to note that simplified estimators are available and accurate in regimes of high SNR and where many trials are available.

### 4.2 Comparison to other methods

We showed that our method strongly outperforms the simple proposal by [20]. Although this competing method was successful on its intended application, here it breaks down since neural activity tends to appear rather deterministically (i.e., spikes occur with very high probability and have low variability in time across trials) for stimuli of high amplitude. This phenomenon is documented in S5, and can be also observed in Figure 2 (see traces in responses to the strongest stimulus). As a consequence, the mean-of-traces estimator of the artifact also contains the neural activity that is being sought, leading to a dramatic failure in detecting spikes, explaining the high false negative rate.

Two other prominent artifact cancellation methods exist, but neither applies directly to our context. The method of [22] considers high-frequency stimulation (5khz). In that context, since action potentials follow a much larger time course than of this very short latency artifact, it is relatively easy to cancel the artifact and recover neural activity by linearly interpolating the recordings whenever stimulation occurs. However, here, as seen in Fig 2, the artifact’s time course can be larger than of spikes (especially at the stimulating electrode). Additionally, the method of [21] has guarantees of success only for latencies greater than 2ms after the onset of stimulus, much larger than the ones addressed here (as small as 0.3 ms). Their 2ms threshold comes from the observation that it is at that time when spikes and artifacts become spectrally separable. However, in our case, at smaller latencies the artifact has a highly transient nature and there is much diversity of artifact shapes (Fig 3) for different electrodes and pulse amplitudes. This immediately excludes the possibility of considering an algorithm based on the spectral differentiation between the spikes and the artifacts in the low-latency context we care about.

### 4.3 Online data analysis, closed-loop experiments

The present findings open a real possibility for the development of closed-loop experiments to achieve selective activation of neurons, [10,45] featuring online data analysis at a much larger scale scale than was previously possible.

We briefly discuss a hypothetical pipeline for a closed loop-experiment, involving four steps: i) visual stimulation and subsequent spike sorting to identify neurons and their Els; ii) single-electrode stimulation scans to map the excitability of those neurons with respect to each of the electrodes in the MEA; iii) additional multi-electrode stimulation to further explore ways to activate cells (optional); and iv) computation of optimal stimulation patterns to match a desired spike train.

Step (iii) might be helpful to enhance combinatorial richness (i.e. the number of ways in which ways neurons can be stimulated) if the available stimulus space resulting from single-electrode stimulation does not lead to a complete selective activation of neurons (in the retina, this will often be the case [44]). There is a caveat, though: allowing for arbitrary stimulation patterns is not possible without further assumptions, since the number of possible amplitude series, i.e., sequences of multi-dimensional stimuli with increasing amplitude, increases exponentially with the number of stimulating electrodes. We propose two solutions: 1) focus on patterns for which there is a clear underlying biophysical interpretation in terms of interactions between the neural tissue and the applied electrical field (e.g., the bipolar and local return stimulation patterns explored here) so that the number of patterns remains bounded, and 2) relax the amplitude series assumption; i.e. allows modes of data collection where recordings are not in response to a sequence of stimulus with increasing strength. This would be possible if artifacts obeyed linear superposition (i.e. the artifact to arbitrary stimulation breaks down into the linear sum of the individual artifacts), since then we would simply need to save the artifacts to single electrode stimulation, and subtract them as required from traces to arbitrary stimuli. In S6 we provide some elementary evidence that supports this linear superposition hypothesis in the simplest, two-electrode stimulation case. However, we stress that further research is required to establish artifact linearity more generally.

### 4.4 Limitations

Here we comment on the current limitations of our method while suggesting some possible extensions.

#### 4.4.1 Beyond the retina: dealing with unavailability of electrical images

We stress the generalizability of our method to neural systems beyond the retina, as we expect that the qualitative characteristics of this artifact, being a general consequence of the electrical interactions between the neural tissue and the MEA [16], are replicable up to different scales that can be accounted for by appropriate changes in the hyperparameters.

In this work we have assumed that the EIs of the spiking neurons are available. At least in the retina, this will normally be the case, as spontaneous firing is ubiquitous among retinal ganglion cells [46]. Thus we can use this spontaneous activity to infer the EIs or other cell properties (e.g. cell type) ‘in the dark’ [47]. If this is not the case, we propose stimulation at low amplitudes so that the elicited cell activity is variable and therefore an initial crude estimate of the artifact can be initialized by the simple mean or median over many repetitions of the same stimulus. Then, after artifact subtraction EIs could be estimated with standard spike sorting approaches.

More generally, this additional EI estimation step could be stated in terms of an outer loop that iterates between EI estimation, given current artifact estimates, and neural activity and artifact estimation given the current EI estimate — that is, our algorithm. Furthermore, we notice the EI estimation step is essentially spike sorting; therefore, there is room for the use of state-of-the-art [48,49] methods to achieve efficient implementations. This outer loop would be especially helpful to enable the online update of the EI in order to counteract the effect of tissue drift, or to correct possible biases in estimates of the EI provided by visual stimulation [50,51], which could lead to problematic changes in EI shape over the course of an experiment. We acknowledge, however, that the implementation of this loop could significantly increase the computational complexity of our algorithm, and deem as an open problem how to achieve a reduction in computational complexity so that online data analysis would still be feasible.

#### 4.4.2 Small spikes: accounting for correlated noise

We assumed that the noise process (*ϵ*) was uncorrelated in time and across electrodes, and had a constant variance. This is certainly an overly crude assumption: noise in recordings does exhibit strong spatiotemporal dependencies [12,52], and methods for properly estimating these structured covariances have been proposed [12,53]. To relax this assumption we can consider an extra, pre-whitening stage in the algorithm, where traces are pre-multiplied by a suitable whitening matrix. This matrix can be estimated by using stimulation-free data (e.g. while obtaining the EIs) as in [12]. The use of a more accurate noise model might be helpful as a means to decrease the signal-to-noise ratio under which the algorithm can operate: here, we discarded neurons whose EI peak strength was smaller than 30 *μ* V (across all electrodes), as the guarantees for accurate spike identification were lost in that case. If this threshold of 30 can be decreased then cells with typically smaller spikes (e.g. retinal midget cells) could be better identified.

#### 4.4.3 Saturation

Amplifier saturation is a common problem in electrical stimulation systems [14,16,[19], and arises when the actual voltage (comprising artifacts and neural activities) exceeds the saturation limit of the stimulation hardware. Although in this work we have considered stimulation regimes that did not lead to saturation, we emphasize that our method would be helpful to deal with saturated traces as well: indeed, in opposition to naive approaches that would lead to no other choice than throwing away entire saturated recordings, our model-based approach enables a more efficient treatment of saturation-corrupted data. We can understand this problem as an example of inference in the context of partially missing observations, for which methods are already available in the GP framework [32].

Finally, notice the above rationale applies not only to saturation, but also to any type of data corruption that could render the recordings at certain electrodes useless.

#### 4.4.4 Automatic detection of failures and post-processing

Since errors cannot be fully avoided, in order to enhance confidence in neural activity estimates provided by the algorithm in the absence of rapid human analysis, we propose to consider diagnostic measures to flag suspicious situations that could be indicative of an algorithmic failure. We consider two measures that arise from a careful analysis of the underlying causes of discrepancies between algorithm and human annotation.

The first comes from the activation curves: at least in the retina, it has been widely documented that these should be smoothly increasing functions of the stimulus strength [25,39]. Therefore, deviations from this expected behavior — e.g., non-smooth activation curves characterized by sudden increases or drops in spiking probability — are indicative of potential problems. For example, the outlier in Fig 6D and many of the false positives in 6C are the result of an incorrectly inferred sudden increase of spiking from one stimulus amplitude to the next (not shown). Moreover, often this sudden increase is ultimately caused by a wrong extrapolation estimate, either with the kernel-based or naive extrapolation estimators. Thus, the application of this simple post-processing criterion (detection of sudden increases in spiking probability) would mark this cell for revised analysis.

The second relates to the residuals, or the difference between observed data and the sum of artifact and neural activity. Cases where those residuals are relatively largecould indicate a failure in detecting spikes, perhaps due to a mismatch between a mis-specified EI and observed data. Indeed, we observed many cases where results were wrong because recordings contained activity that did not match any of the available templates (not shown). In such cases it is hard even for a human to make a judgment, as he or she has to carefully decide whether the observed activity corresponds to an available inaccurate EI or rather, to a truly spiking neuron that was not identified during the EI creation stage. We have reported these as errors, but we highlight they were propagated from the previous spike sorting stage. Therefore, methods to quantify the per-neuron credibility of the templates, such as those developed in [54], are of crucial importance here to complement the above residual criterion.

In either case, the diagnostic measures can be implemented as an automatic procedure based on goodness-of-fit statistics (e.g. the deviance [55]), or even simpler quantities (e.g. an abrupt increase in firing probability between two consecutive values). Moreover, we have showed in related work [56] that these automatic diagnostics can be implemented in a further post-processing stage, where the artifact is locally re-sampled or interpolated from the Gaussian model if a possible error has been diagnosed.

#### 4.4.5 Larger and denser arrays, different time scales

In this work, the computationally limiting factor is *E*, the number of electrodes, as this dominates the (cubic) computational time of the GP inference steps. Recent advances in the scalable GP literature [57–59] should be useful for extending our methods to even larger arrays as needed; we plan to pursue these extensions in future work.

Finally, we also note that an extension to denser arrays (e.g. [60]) is immediately available within our framework: indeed, preliminary results with denser arrays (30*μm* spacing between electrodes, not shown) revealed that due to the increased proximity between the stimulating electrode and its neighboring electrodes, those electrodes also possessed large artifacts and were subject to the effect of breakpoints. Then, we can proceed exactly as we did in 2.5.2 for local return, by considering different models for the stimulating electrode and its neighbors.

## 5 Conclusion

We have developed a method to automate spike sorting in electrical stimulation experiments using large MEAs, where artifacts are a concern. We believe our developments will be useful to enable closed-loop neural stimulation at a much larger scale than was previously possible, and to enhance the ability to actively control neural activity. Also, our algorithm has the potential to constitute an important computational substrate for the development of future neural prostheses, particularly epi-retinal prostheses. We have made available, in the first author’s website, MATLAB code that contains an example applying the algorithm to process one of the datasets analyzed in this paper.

## 6 Acknowledgments

LEG received funding from NIH Grant 1F32EY025120. EJC received funding from NEI Grant EY021271. LP received funding by NSF BIGDATA IIS 1546296. JPC received funding from Sloan Foundation and the McKnight Foundation fellowships. PH received funding from Polish National Science Centre grant DEC-2013/10/M/NZ4/00268. We thank David Blei Nishal Shah for useful discussions, the anonimous reviewers for helpful feedback, Frederick Kellison-Linn and Victoria Fan for manual data analysis, Georges Goetz and Pulkit Tandon for computational assistance, and Mariano Gabitto and Ella Batty for their comments on the manuscript.

## 7 Author Contributions

Conceived and designed the methods/experiments: GEM, LP, JPC, LEG, EJC. Performed the experiments: LEG, SM. Analyzed the data: GEM, LG, SM, LP. Contributed reagents/materials/analysis tools: AL, PH, EJC. Wrote the paper: GEM, LP, EJC, LEG, JPC.

## S1 Text

### Experimental procedures

All electrophysiology data were recorded from primate retinas isolated and mounted on an array of extracellular electrodes as described in previously published literature [39]. Eyes were obtained from terminally anesthetized macaque monkeys (Macaca species, either sex) used for experiments in other labs, in accordance with IACUC guidelines for the care and use of animals. After enucleation, the eyes were hemisected and the vitreous humor was removed. The hemisected eye cups containing the retinas were stored in oxygenated bicarbonate-buffered Ames solution (Sigma) at room temperature during transport (up to 2 hours) back to the lab. Patches of intact retina 3mm in diameter were isolated and placed retinal ganglion cell-side down on a 512-electrode MEA. Throughout the experiments, retinas were superfused with oxygenated bicarbonate-buffered Ames solution at 35°C.

In all experiments the raw voltage signals from each electrode were amplified, filtered, and multiplexed with custom circuitry [16,61]. Electrodes had diameters of 10-15 *μ*m and were separated by 60 *μ*m. Data were acquired at 20 kHz on all electrodes and bandpass filtered between 43 and 5000 Hz. Charge-balanced, triphasic current pulses with relative amplitudes of 2:-3:1 and phase widths of 50 *μs* were applied to each electrode, and reported current amplitudes correspond to the charge of the second, cathodal, phase. A platinum ground wire circling the perfusion chamber served as a distant ground in all one-electrode stimulation experiments. In some experiments, a 1 mM tetrodotoxin (TTX) solution in Ames solution was perfused into the retina to inhibit all action potentials in order to directly measure the stimulus artifact in a retinal preparation.

#### Obtaining the EIs

Retinal ganglion cells (RGCs) were identified in the absence of electrical stimulation using previously described spike sorting techniques [27] and classified into types based on how they respond to a visual white noise stimulus projected onto the retina [62,63]. For each RGC, thousands of voltage waveforms were averaged on all electrodes, resulting in a spatiotemporal voltage signature specific to that RGC. These signatures are used as templates in our sorting algorithm.

#### Estimation of mean

Regarding the mean parameter of the artifact kernels, *μ*, we follow the standard in the applied statistics community: *μ* is a centering parameter and all the non-random aspects of data should be captured by it. In our case this component is given by what we call the switching artifact, a waveform *A*_0_ = *A*_0_(*e*, *t*) that is present regardless of the amplitude of stimulation. We estimate 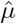 by taking the mean of recordings at the lowest amplitude of stimulation (see S1 Fig for details on the characteristics of the switching artifact, and to see the effect of this mean-subtraction stage on recordings).

### Dataset details

#### Real data

##### Population statistics, data selection

In total, we analyzed 4,045 amplitude series coming from thirteen retinal preparations, giving rise to 1,713,223 trials. These amplitude series are the ones for which reliable human curated data was available. The human analysis of these datasets was required by various previous research projects (see for example [28,41,[44], where the human analysis procedure is explained). In Table 1 we specify details of the thirteen retinal preparations for which human annotation (HA) was available. In some preparations (e.g. 2012-09-24) there is human annotated data from multiple stimulation modalities. Also, in Table 2 we specify the population statistics of activation, both in terms of spikes and activation in amplitude series.

For each preparation and stimulus modality, there were characteristic numbers of stimulation patterns and neurons being analyzed. Usually, given a stimulating electrode, human annotation was available for only one, or at most a few neurons (e.g. two or three). However, we considered the totality of EIs of neurons that had strong enough signals (overall EI peak strength greater than 30 *μ*V and 8*μ*V at at least one stimulating electrode) but restricted performance computations to the subsets of neurons for which human annotation was available.

##### Bundle detection

Importantly, we restricted our analysis to the stimulation amplitudes that did not lead to gross contamination of recordings due to the activation of entire axonal bundles in the retina (for a recent account of this pervasive phenomenon see [44]), as this would lead to a situation that is not accounted for by our model. For each amplitude series with available human annotation, we determined the maximum amplitude of stimulation that did not lead to activation of a bundle by looking for ‘hot’ electrodes, distant from the stimulating one, exhibiting high temporal variance in the artifact (here, for simplicity the artifact was estimated by the simple average over traces). Then, we did not consider any amplitude of stimulation beyond the onset of axonal bundle activation, the first amplitude where we identified such hot electrodes. We found that a robust method for estimating this threshold (equivalently, the presence of hot electrodes) was based on a Kolmogorov-Smirnov goodness-of-fit test on the empirical distribution of the (log) temporal variances of the artifact on distant electrodes, with the Gaussianity null hypothesis. The appearance of hot electrodes created a new mode in the distribution, leading to a violation of the normality assumption. We found that by setting the cut-off *p*-value for this test as 10^−12^ we achieved the best match with axonal bundle activation onsets estimated by human experts (not shown).

##### Refractory period

We considered time windows of 2*ms* (*T* = 40, at a 20khz sampling rate), which is smaller than the usual refractory periods of retinal ganglion cells [64,65], and which in practice did not lead to multiple neural events for the same neuron on the same trial. Also, spikes were sought in the interval [0.35,1.35] ms following the onset of the 150 *μs* triphasic stimulus. This interval encompasses the range were most of the artifact variation occurs; that is, where non trivial artifact cancellation methods are required.

##### Parallel analysis

For the analysis in Fig 6I we reported times and their variability — the experiment was repeated ten times — for the analysis of the eight single-electrode scans for which for which some human-curated data was available (see Table 1 S1 Text for details on those retinal preparations). These experiments were done on an Intel Xeon E5-2695V2 12C/24T 2.4Ghz 8.0GT/s 30mb CPU, with 20 threads running in parallel.

**Table 1.** Details of the retinal preparations analyzed for each type of stimulation: *Single Electrode* (S.E.), *Bipolar*(B.), *Local Return* (L.R.) and *Arbitrary* (A). stimulation

| Preparation ID | Type | #Neurons in preparation | #Neurons with HA | #Trials | #Amplitude series with HA | #Trials per stimulus |
| --- | --- | --- | --- | --- | --- | --- |
| 2012-09-24-3 | S.E. | 559 | 36 | 400,805 | 333 | 51 |
| 2014-09-10-0 | S.E. | 378 | 5 | 40,802 | 33 | 48 |
| 2014-11-05-3 | S.E. | 322 | 19 | 37,940 | 72 | 21 |
| 2014-11-05-8 | S.E. | 277 | 19 | 37,644 | 71 | 21 |
| 2014-11-24-2 | S.E. | 439 | 11 | 36,078 | 94 | 21 |
| 2015-04-09-2 | S.E. | 252 | 6 | 31,775 | 49 | 25 |
| 2015-04-14-0 | S.E. | 623 | 20 | 86,655 | 138 | 25 |
| 2015-05-27-0 | S.E. | 332 | 8 | 30,368 | 38 | 25 |
| Total | S.E. | 3,182 | 124 | 702,067 | 828 | n.a. |
| 2012-09-24-3 | B. | 559 | 34 | 187,612 | 248 | 30 |
| 2012-09-27-4 | B. | 482 | 17 | 170,787 | 184 | 50 |
| 2014-11-24-2 | B. | 439 | 9 | 32,395 | 70 | 30 |
| 2015-03-09-0 | B. | 409 | 6 | 67,332 | 58 | 42 |
| 2015-04-09-2 | B. | 252 | 7 | 83,143 | 79 | 42 |
| 2015-05-27-0 | B. | 332 | 8 | 65,023 | 42 | 50 |
| Total | B. | 2,473 | 81 | 606,292 | 681 | n.a. |
| 2014-11-24-2 | L.R. | 439 | 14 | 43,822 | 104 | 21 |
| 2015-04-09-2 | L.R. | 252 | 4 | 15,624 | 27 | 25 |
| 2015-04-09-3 | L.R. | 569 | 2 | 9,575 | 15 | 25 |
| 2015-04-14-0 | L.R. | 623 | 25 | 60,597 | 98 | 25 |
| 2015-09-23-2 | L.R. | 686 | 28 | 28,574 | 56 | 25 |
| Total | L.R. | 2,569 | 73 | 158,192 | 300 | n.a. |
| 2015-05-27-0 | A. | 332 | 4 | 246,672 | 2,236 | 10 |
| Total | A. | 332 | 4 | 246,672 | 2,236 | n.a. |
| Grand Total | All | 4443 | 282 | 1,713,223 | 4,045 | n.a. |

#### Simulated data

Simulated data was created by artificially adding neural activity to TTX recordings, in an attempt to faithful mimic the phenomena observed in the real case [26,39]. Specifically, we considered 83 neurons (the largest subset of the ones targeted in the single-electrode real data analysis so that their EIs did not heavily overlap) and recordings to 380 stimulating electrodes (one at a time) in a TTX experiment with *n*_*j*_ = 6 trials to *J* = 35 different stimuli between 0.1 and 3.5*μA*. Then, given a single stimulating electrode we sampled activation curves for all the neurons whose EI at the stimulating electrode was strong enough, indicating proximity. Activation curves were parametrized by their thresholds, chosen uniformly in the stimulation range, and their steepness, also sampled uniformly. Spikes of those neurons were then sampled from these activation curves with latencies chosen so they would match the human spike sorting results (summarized in S4 Fig) in the following two aspects: 1) they had same median latency as a function of the distance between the neuron and stimulating electrodes (spiking of nearby neurons has shorter latency) and 2) they had same variance in spike latency as a function of spike probability (in the steady spiking regimes, where the probability of firing is high, latencies are much less variable). Also, to obtain better estimates of false positive rates, we fed the algorithm with ‘dummy’ neurons (three per amplitude series, with EIs chosen at random from the available set of remaining neurons) with no spiking at all.

**Table 2.** Population frequency of activation events, for the trial-by-trial and amplitude-series based analysis.

| Type of stimulation | Trial based |  | Amplitude series based |  |
| --- | --- | --- | --- | --- |
|  | #Trials | #Trials with spikes | #Amplitude series | #Amplitude series with activation |
| Single Electrode | 702,067 | 15,830 | 828 | 36 |
| Bipolar | 606,292 | 26,535 | 681 | 100 |
| Local Return | 158,192 | 3,564 | 300 | 11 |
| Arbitrary | 246,672 | 16,219 | 2,236 | 293 |
| All | 1,713,223 | 62,148 | 4,045 | 440 |

All the reported results involving simulations are based on 5000 samples of amplitude series following the above procedure.

## S2 Text

Here we review the main algebraic properties, summarized in [34], that we implement to achieve fast kernel computations. In all of the below, *K*_*d*_, (*d* = 1,…, *D*) are square invertible matrices with dimensions *n*_*d*_.

**Property 0.** Associativity. The Kronecker product is associative.

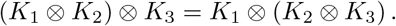

**Property 1.** Inversion of the Kronecker product. The inverse of a Kronecker product equals the product of their inverses: 
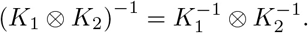

**Property 2.** Kronecker product eigen-decomposition. If

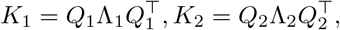

then

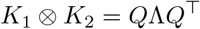

where

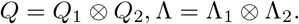

In other words, the eigen-decomposition of a Kronecker product corresponds to the product of their eigen-decompositions.

**Property 3.** Trace of a Kronecker product. The trace of a Kronecker product is the product of the individual traces:

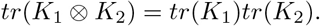

**Property 4.** Log determinant of the Kronecker product. The log determinant of the Kronecker product is a weighted sum of the individual log determinants, and the weights are the dimensions:

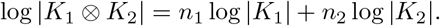

**Property 5.** Matrix product between a Kronecker product and a vector. Let *v* be a 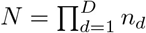 dimensional vector, with each *n*_*d*_ of comparable magnitude. Then

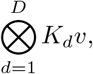

can be computed efficiently in *O*(*DN*^(*D*+1)/*D*^) space and time. For implementation details see algorithm 2 in [33], and our code.

### S1 Fig

**Fig. 1.**
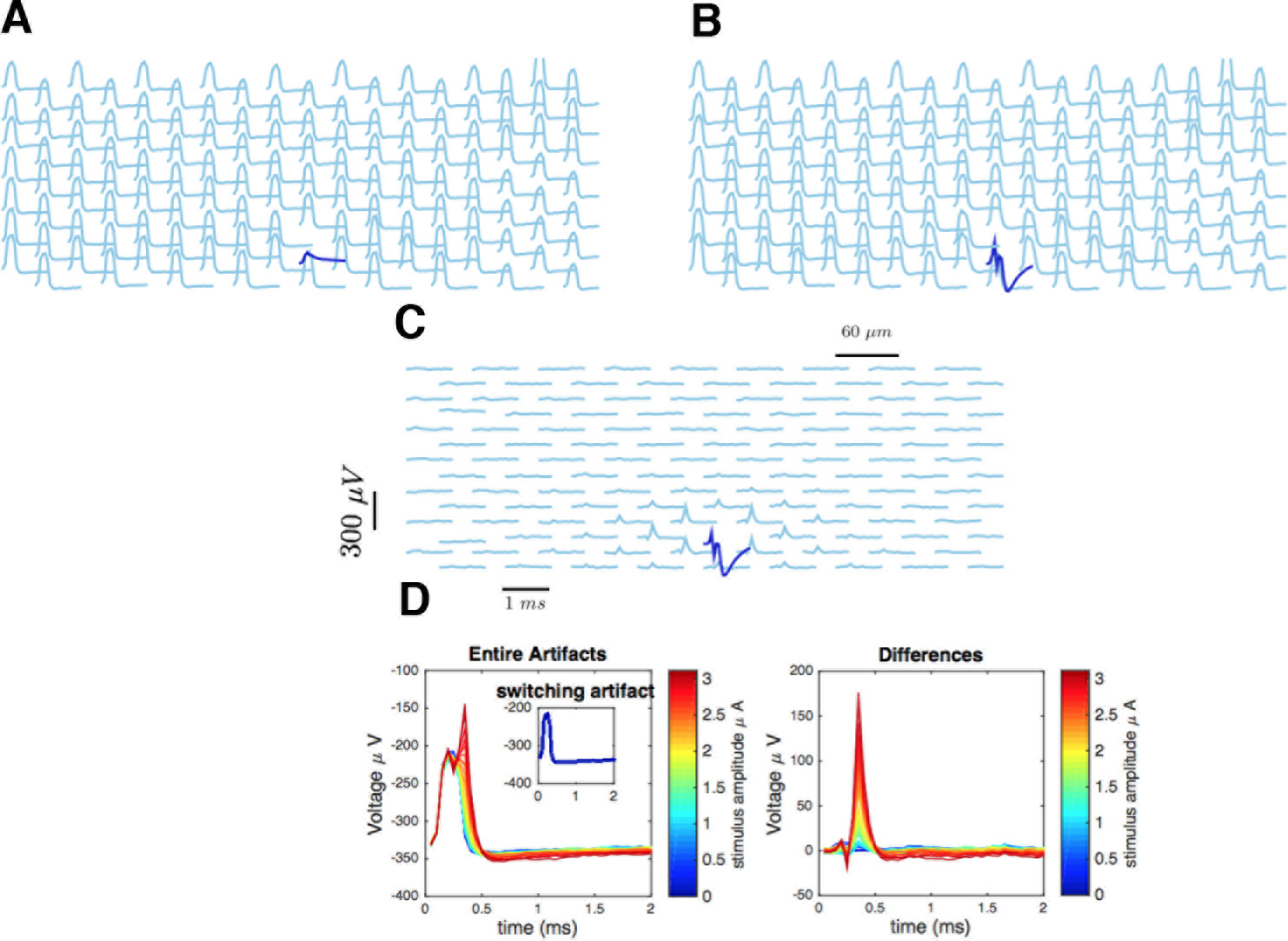
**A** Raw artifact traces at the smallest amplitude of stimulation (0.1 *μA*), considered an estimate of *μ*, the switching artifact. **B** Raw artifact traces at 0.99 *μA* of stimulus. **C** Difference. Notice that the main text refers to this already mean-subtracted artifact. **D**) *Left*: Raw artifact at all different stimuli for a non-stimulating electrode (inset, switching artifact). *Right*: Differences.

### S2 Fig

**Fig. 2.**
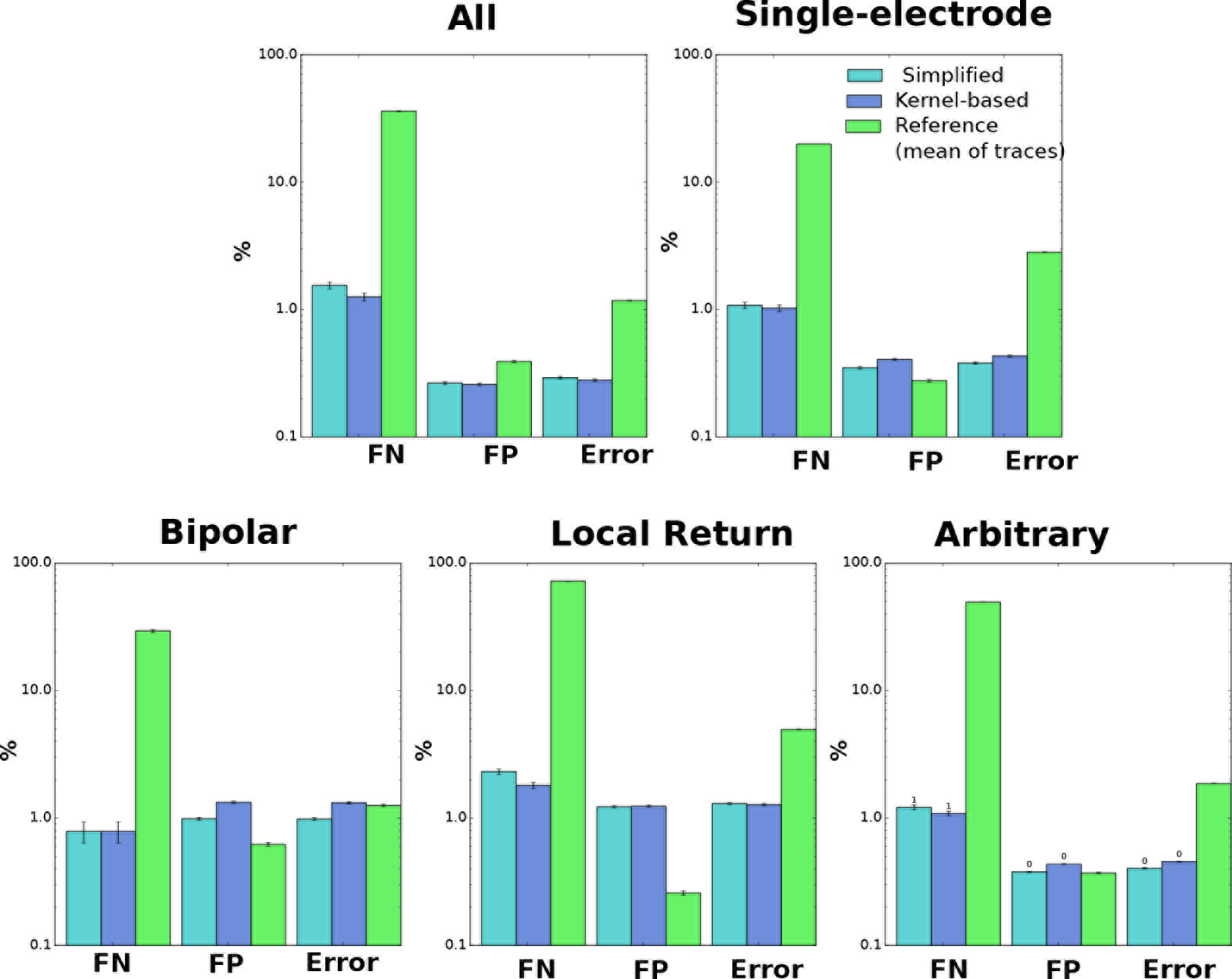
**Population Results** (log scale) including the mean-of-traces estimator proposed in [20] and our simplified estimator. These results complement Figure 6A, by reporting differences by type of estimator, and also by reporting total errors.

### S3 Fig

**Fig. 3.**
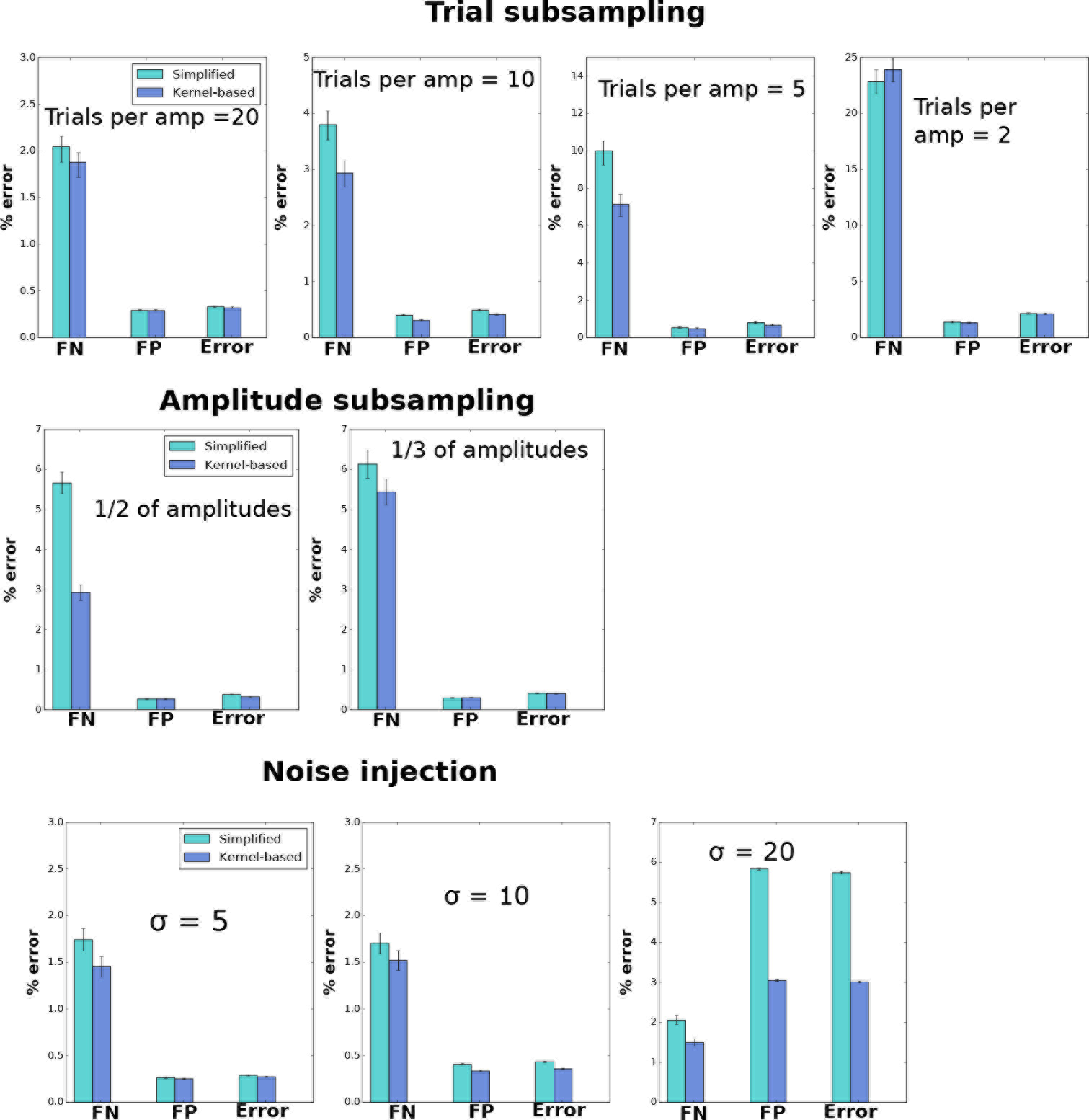
Comparison of simplified and kernel-based estimator in the analysis of perturbations to real data. These results complement Figure 6B, by reporting false positive and negative rates at different conditions for trial subsampling (top), amplitude subsampling (middle) and noise injection (bottom). Only for single electrode stimulation. Notice that for trial sub-sampling and noise injection, results may vary from one experiment to another.

### S4 Fig

**Fig. 4.**
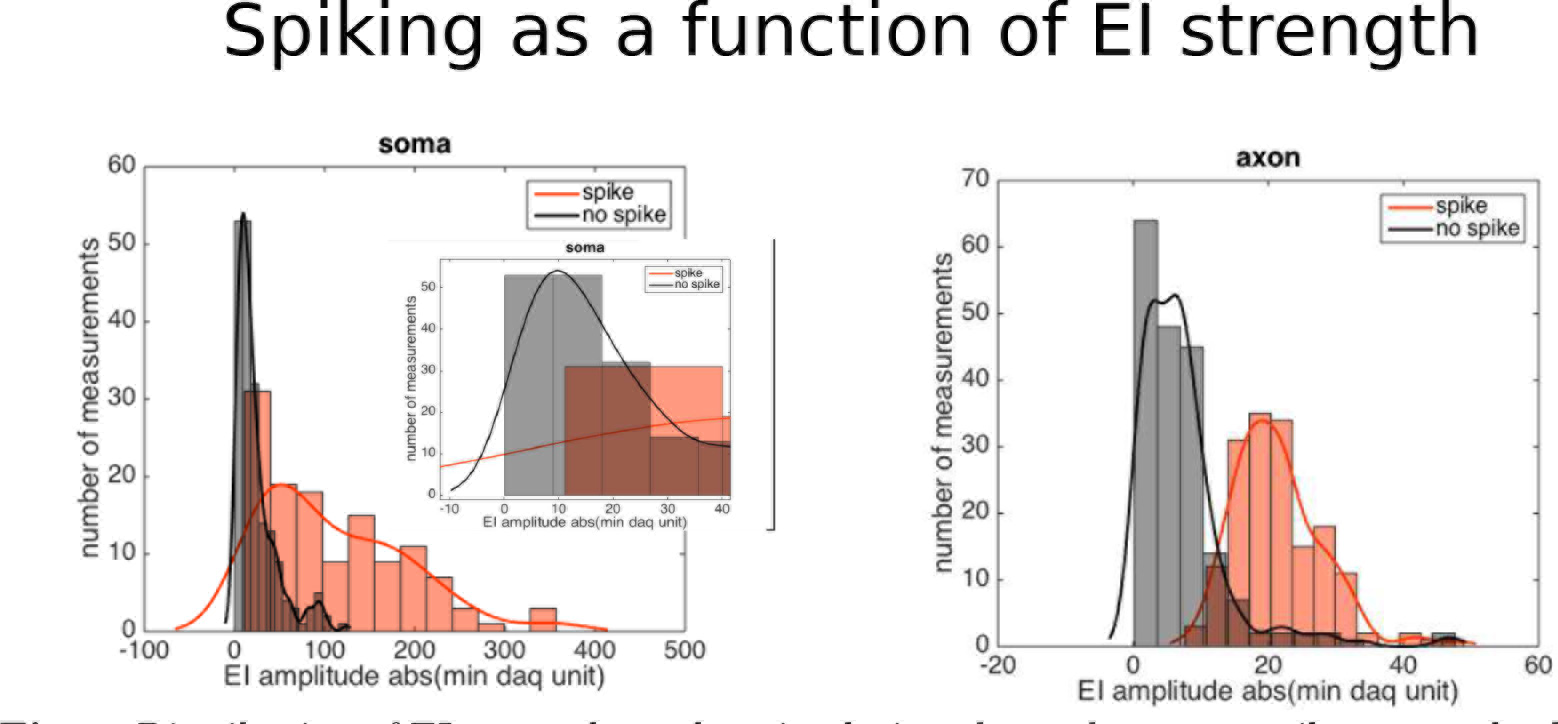
Distribution of EI strength on the stimulating electrode among spike events, both for somatic and axonal (distant) stimulation. For somatic stimulation inset corresponds to a zoom to smallest voltages. For EI peak strengths smaller than 10*μV* spike is not observed (based on manual analysis).

### S5 Fig

**Fig. 5.**
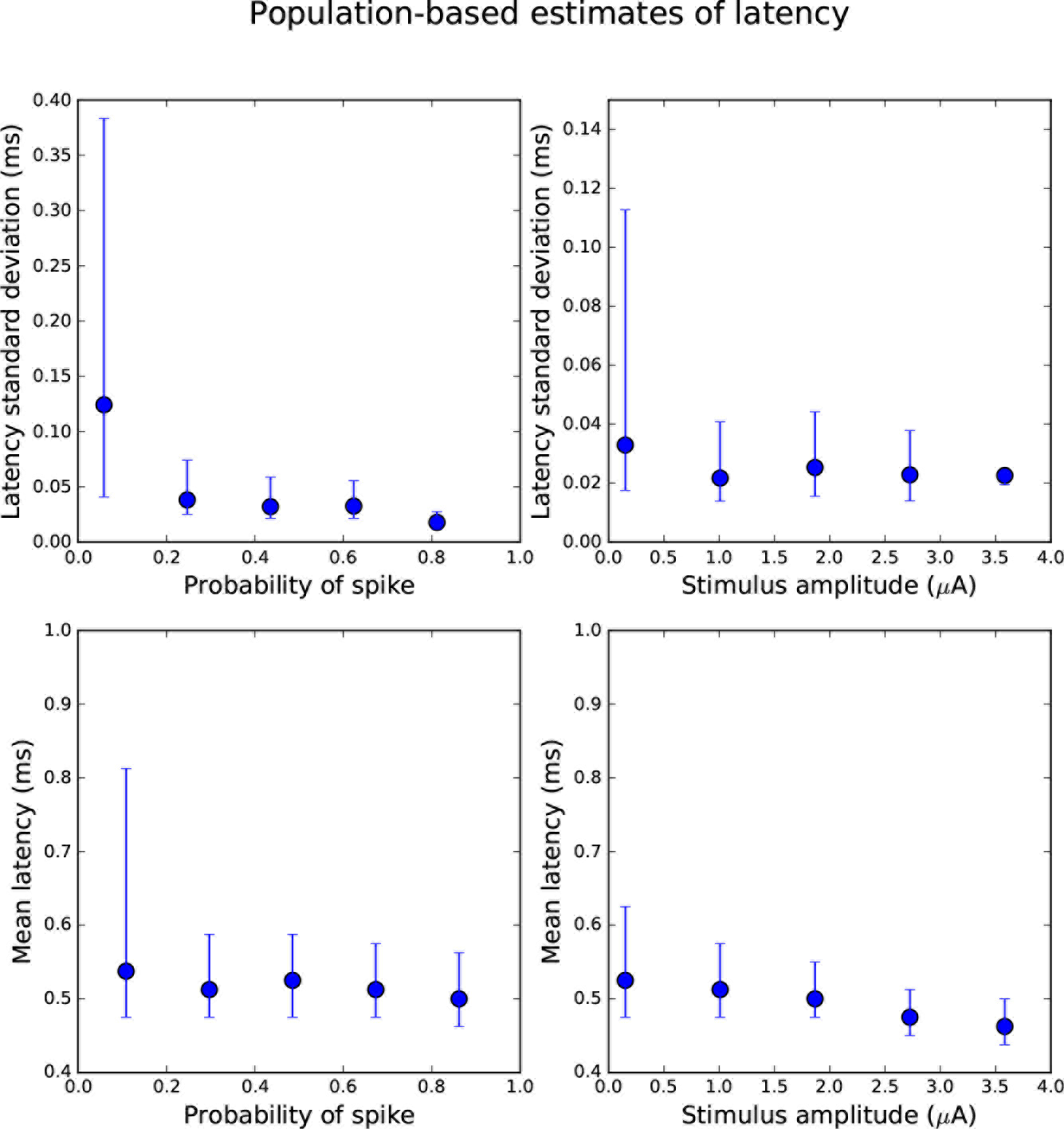
Population based estimates of the mean (top) and standard deviation (bottom) of spike latency, as a function of probability of spiking (left) and stimulus amplitude (right). This supports the observation that when activation is reached (high probability of spike) variability of latencies reaches its minimum.

### S6 Fig

**Fig 6.**
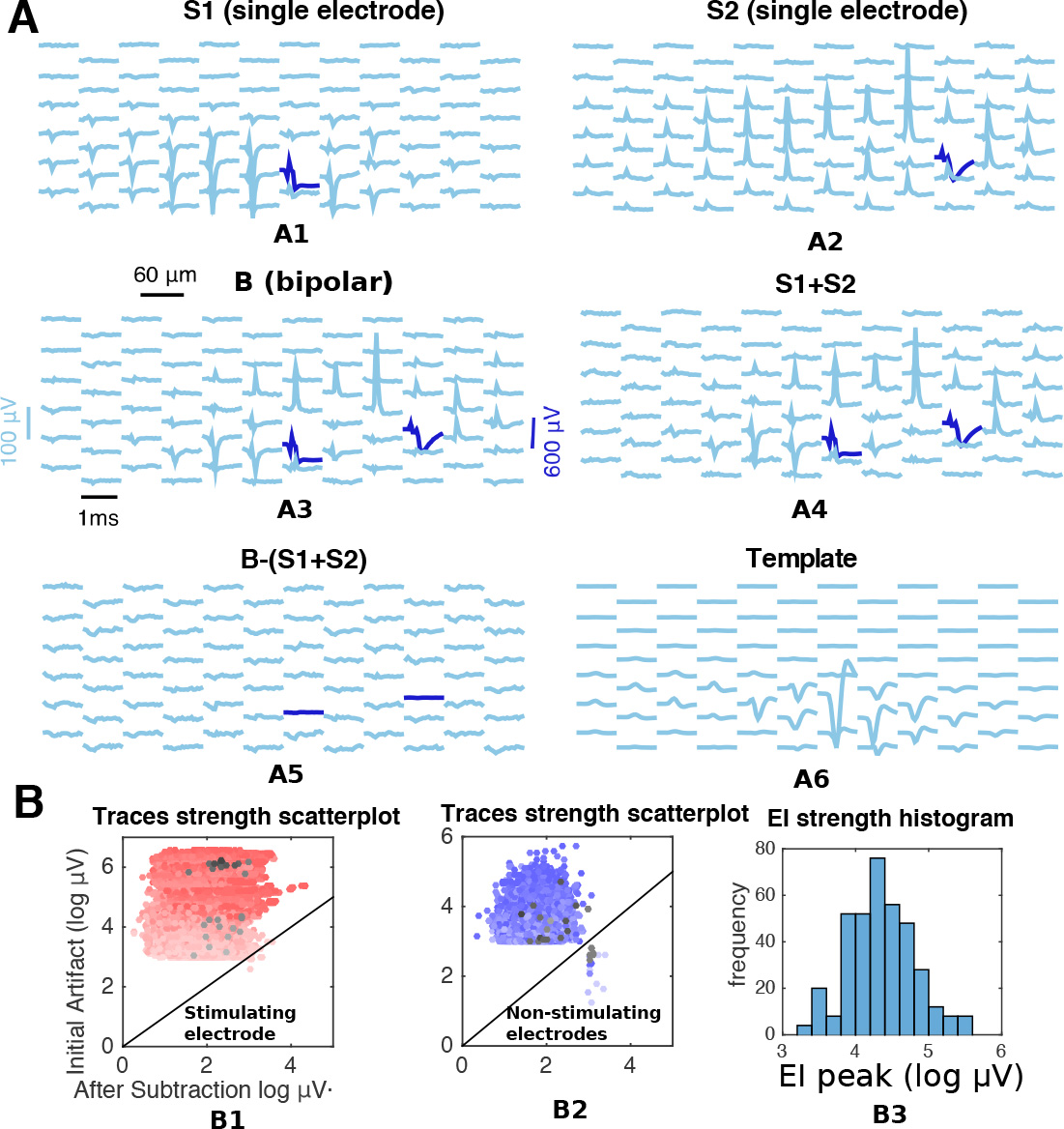
The linear superposition of artifacts provides a reasonable phenomenological model for two electrode stimulation. Observations are based on a single retinal preparation (TTX). **A**) example of observed linearity: *A1*-*A2*) artifacts for single electrode stimulation at two different stimulating electrodes with same strength (3.1 *μ* A) and opposite polarities. *A3*) corresponding two-electrode stimulation. *A4*) sum of *A1*) and *A2*). *A5*) difference between *A3*) and *A4*). *A6*) for reference, the EI of a typical neuron in shown in the same scale. **B**) population-based generalization of the finding in **A**) from thousands of stimulating electrode pairs, collapsing stimulating amplitudes and electrodes. *B1*-*B2*) scatterplots of the maximum strength (over electrodes and time) of two-electrode stimulation artifacts at different stimulus strengths (strength of the color) before and after subtracting the sum of single electrode artifacts. Points in the gray-scale are the ones shown in *A*). *B3* histogram of log peak EI of neurons in the array. In the light of *B3, B1,B2* show in the vast majority of artifacts of magnitude comparable with than of EI ( 99% of points above the diagonal and outside the log-strength 2.5 *μV* boxes in *B1,B2*) subtracting the linear sum of individual artifacts is a sensible choice as it decreases its strength.

